# Natural genetic variation spans the qualitative phenotypic space of a neuronal identity endophenotype

**DOI:** 10.64898/2026.06.05.730093

**Authors:** Dorian Natario, Sonya Galant, Joseph R. Bunker, Benjamin Mormann, Francesca Viscido, Sophie Lin, Rossella E. Ciliberti, Deepshe Dewett, Faredin Alejevski, Laurent Théodore, Annalise B. Paaby, Jens Rister, Daniel Vasiliauskas

## Abstract

Understanding the genetic basis of natural phenotypic variation remains challenging, particularly for complex traits. However, causal genetic variants and their effects may be more accessible through endophenotypes, intermediate traits along the trajectory from genotype to complex organismal phenotype. Here, we investigate natural variation in R8 photoreceptors of the *Drosophila* visual system, using Rhodopsin 5 (Rh5) and Rhodopsin 6 (Rh6) expression as a readout of their subtype identities. Simple rules governing this expression allowed us to predict eight qualitatively distinct phenotypes. We observed deviations from the typical R8 subtype pattern in individual flies extracted from the field and in wild-derived inbred lines. Remarkably, we found all eight predicted phenotypes, including shifts in the proportions of Rh5- and Rh6-expressing R8s, Rh5/Rh6 co-expression, and loss of detectable Rh5 or Rh6 expression. We identified causal coding and regulatory variants in *sevenless*, *Rh5*, *Rh6*, and *melted*, affecting different levels of the well-characterised developmental program. Our findings demonstrate that standing genetic variation in natural populations spans the full qualitative phenotypic space of a stereotyped neuronal system, suggesting that substantial deviations, including those associated with likely loss-of-function alleles, can be tolerated in nature.

## INTRODUCTION

Understanding how complex phenotypes evolve requires insight into the nature of genetic variation underlying quantitative traits. However, identifying the mutations responsible for naturally occurring phenotypic differences remains challenging, particularly for complex traits influenced by many variants of small effect^1,2^. A powerful approach to this problem is to investigate natural variation in endophenotypes, intermediate molecular, cellular, or developmental phenotypes that link genetic variation to complex organismal traits^3^. Because they are more proximal to gene action, endophenotypes can reveal functional effects of genetic variants that may be difficult to detect in higher-order traits more closely related to fitness. A well-defined endophenotype can also make it possible to directly assess the range of phenotypic outcomes produced by standing genetic variation. Here, we examine natural variation in the neuronal differentiation of R8 photoreceptors in the *Drosophila* retina, an endophenotype for vision.

The retina comprises approximately 800 unit eyes, called ommatidia (Fig. 1A), each containing eight photoreceptor neurons (R1–R8) arranged in a stereotyped pattern (reviewed in Ref. ^4^). The six outer photoreceptors, R1–R6, mediate dim-light vision and motion detection, and are functionally analogous to vertebrate rods. The two inner photoreceptors, R7 and R8, are involved in colour vision, are functionally analogous to vertebrate cones, and are stacked along the same light path, with R7 above R8.

**Figure 1.**
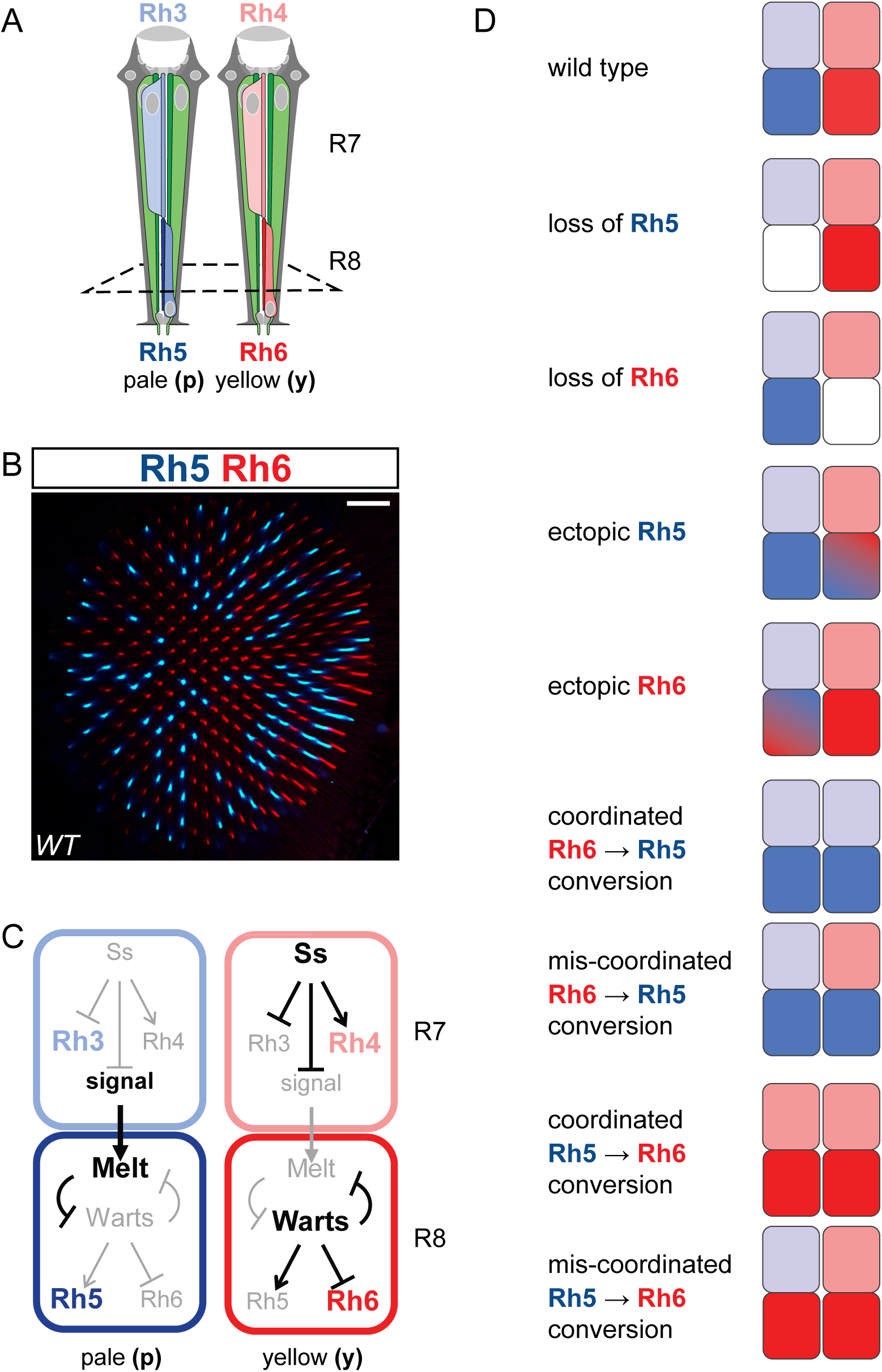
R7–R8 Rhodopsin patterns and categories of Rh5/Rh6 expression change. A. Schematic side views of yellow (left) and pale (right) ommatidia. The lens is positioned at the top; grey cells indicate non-neuronal cells, and coloured cells indicate photoreceptors. R1–R6 photoreceptors (green) express Rh1. Colour-vision photoreceptors (reds and blues) are arranged vertically: the upper photoreceptor, R7, expresses Rh3 (light blue) in pale ommatidia or Rh4 (light red) in yellow ommatidia, whereas the lower photoreceptor, R8, expresses Rh5 (dark blue) in pale ommatidia or Rh6 (dark red) in yellow ommatidia. Adapted from Ref. ^4^. B. A single confocal optical section from a wild-caught fly retina, stained with anti-Rh5 and anti-Rh6 antibodies, shows a wild-type Rh5 (blue) and Rh6 (red) expression pattern in the central region of the retina. The section is transverse to the ommatidial axis at the R8 level (as indicated by the dashed line in (A)). Rh5- and Rh6-expressing R8 photoreceptors (pR8 and yR8) are distributed stochastically in an approximately 1:2 ratio, with no Rh5/Rh6 co-expression. Scale bar, 20 µm. C. Illustration of key steps in R7 and R8 subtype specification in pale (p) and yellow (y) ommatidia. See Introduction for details. D. Schematic showing the eight predicted categories of qualitative Rh5/Rh6 expression changes in R8 (left), encompassing all qualitatively distinct outcomes. Conversion phenotypes are subcategorized based on whether they are coordinated with R7 subtype identity. Colours indicate Rh5 (dark blue) and Rh6 (dark red) expression in R8 and Rh3 (light blue) and Rh4 (light red) expression in R7. Graded blue/red indicates Rh5/Rh6 co-expression. See Introduction for details.

Photoreceptors’ spectral sensitivities, a central aspect of their functional identities, depend largely on the Rhodopsins they express. These light-sensitive proteins absorb photons to initiate phototransduction and are critical for colour discrimination, spatial navigation, motion detection, and circadian synchronisation^5–13^. Their expression provides a direct cellular readout of photoreceptor subtype identity and can be scored both as discrete single-cell states and as quantitative whole-retina phenotypes. Moreover, Rhodopsin expression is controlled by a developmental program whose logic is sufficiently well understood to relate natural variation in expression to defined steps of photoreceptor differentiation. These features make Rhodopsin expression particularly well suited for examining natural variation in a well-defined neuronal differentiation readout. However, variation in Rhodopsin expression patterns, both within and among fly species, has been examined in only a few studies^14–19^.

In the main part of the retina, ommatidia are predominantly of two types, “pale” (p) and “yellow” (y) (Fig. 1A, reviewed in Ref. ^4^). These are distinguished by Rhodopsin expression in the inner photoreceptors. In p ommatidia, R7 (pR7) expresses UV-sensitive Rh3, and R8 (pR8) expresses blue-sensitive Rh5. In y ommatidia, R7 (yR7) expresses a different UV-sensitive Rh4, and R8 (yR8) expresses green-sensitive Rh6. p and y ommatidia are distributed stochastically in an approximate 1:2 ratio (p:y) (Fig. 1B). Each photoreceptor typically expresses only one Rhodopsin gene, except for yR7s in the dorsal retina, which express Rh4 together with varying levels of Rh3.

After R7 and R8 photoreceptors have been specified, their p and y subtypes are determined by a well-characterised developmental program (Fig. 1C) that proceeds as a multistep cascade. In R7 photoreceptors, the transcription factor Spineless (Ss) is expressed in a stochastic on/off pattern^20–22^, with expression in roughly two-thirds of R7s. Ss expression represses the default Rh3-expressing pR7 identity and specifies the Rh4-expressing yR7 identity^20,23^. pR7 cells signal to their underlying R8 partners through the TGFβ pathway^24^. In R8, the Hippo pathway is active by default^25–29^. The pR7-derived signal upregulates the *melted* (*melt*) gene, which encoded a pleckstrin homology-domain protein, and represses *warts* gene, which encodes the core kinase of the Hippo pathway, thereby turning Hippo signalling OFF. Mutual transcriptional repression involving Melted and Warts forms a bistable switch that stabilises either the default yR8, Rh6-expressing state or, after pR7 signalling, the alternative pR8, Rh5-expressing state, thereby committing the cell to one of these two fates^25–29^. As a result, each R8 expresses either Rh5 or Rh6, but not both. In the ageing adult, additional active molecular mechanisms maintain this mutually exclusive Rh5/Rh6 expression pattern, and their disruption can give rise to age-dependent phenotypes^30^. The mutually exclusive Rh5/Rh6 expression pattern therefore provides a sensitive readout of the R7–R8 differentiation and maintenance cascade, reflecting both direct effects on R8 fate and, via the coordinating signal, upstream changes in R7 subtype identity. In the experiments described below, we exploit this readout in aged flies to investigate natural genetic variants that may perturb this developmental and maintenance network.

Because Rh5/Rh6 expression in R8 is a mutually exclusive binary choice, six qualitative phenotypic changes can be predicted: loss of Rh5 or Rh6 expression; ectopic gain of Rh5 or Rh6 expression in the wrong R8 subtype, resulting in Rh5/Rh6 co-expression in yR8 or pR8, respectively; and conversion from Rh5 to Rh6 or from Rh6 to Rh5 (Fig. 1D). These six categories encompass all possible qualitatively distinct changes in Rh5/Rh6 expression in R8 photoreceptors (quantitative differences in Rh5 and Rh6 expression levels within individual cells are not considered here). The conversion phenotypes can be further subdivided into those coordinated with R7 subtype changes and those occurring independently of R7 identity, yielding a complete set of eight theoretically predicted qualitative phenotypes.

Seven of these eight predicted phenotypes have been reported in laboratory-generated mutants that perturb the differentiation and maintenance program (see refs.^20–30^ for examples). In these mutants, generally only a fraction of R8s is affected; this fraction is usually consistent for a given genotype and can be quantified precisely and reproducibly. Thus, phenotypes that are qualitative at the cellular level are quantitative at the whole-eye level, ranging from mild to strong. In particular, incomplete conversion phenotypes manifest as shifts in the overall p:y R8 ratio, reflecting changes in the relative frequencies of Rh5- and Rh6-expressing cells.

To begin exploring natural variation in Rh5/Rh6 expression, we examined wild *Drosophila melanogaster* collected from natural populations. The marked diversity of Rh5/Rh6 phenotypes we observed among individuals suggested that naturally segregating genetic differences might contribute to this variation.

To test this under genetically reproducible conditions, we took advantage of the DGRP2 (*Drosophila* Genetic Reference Panel release 2), a collection of 205 inbred fly lines each derived from a single wild female^31–33^. The genomes of these lines have been sequenced, and more than 4.5 million naturally occurring molecular variants have been computationally identified^31,32^. The collection has been successfully used to investigate the genetic architecture of numerous quantitative traits^33^; however, only a few studies have identified the actual causal genetic changes. A relevant prior study is the work by Anderson et al.^16^, which screened for natural variants in the *ss* locus that affect the R7 p:y ratio. That work identified a single-base insertion allele of *ss*, called *sin,* which increases the p:y R7 ratio.

Surprisingly, we found all eight predicted Rh5/Rh6 phenotypes among inbred lines. We identified specific coding and regulatory variants responsible for several of these phenotypes, linking natural variation to different components of the photoreceptor differentiation and maintenance program. The breadth of variation in this normally stereotyped neuronal system suggests that even substantial deviations, including those associated with likely loss-of-function alleles, can be tolerated in nature.

## RESULTS

### Variation of Rh5/Rh6 expression in wild flies

To investigate variation in Rh5/Rh6 expression patterns in R8 photoreceptors in wild populations, we captured flies from multiple locations in France and the UK (see Materials and Methods for details). Retinas of either the captured flies themselves or one F1 progeny from each captured female (i.e., flies resulting from mating in the wild) were stained with specific antibodies to reveal Rh5 and Rh6 expression, and in some samples, Rh4 expression was also examined. Prior to staining, flies were aged for over two weeks to detect maintenance phenotypes in addition to the specification phenotypes. To quantify the phenotypes, we counted the number of R8 photoreceptors expressing Rh5 or Rh6, those co-expressing both Rhodopsins, and those expressing neither. Percentages of each R8 class were then calculated (Fig. 2A, Supplementary Data 1).

**Figure 2.**
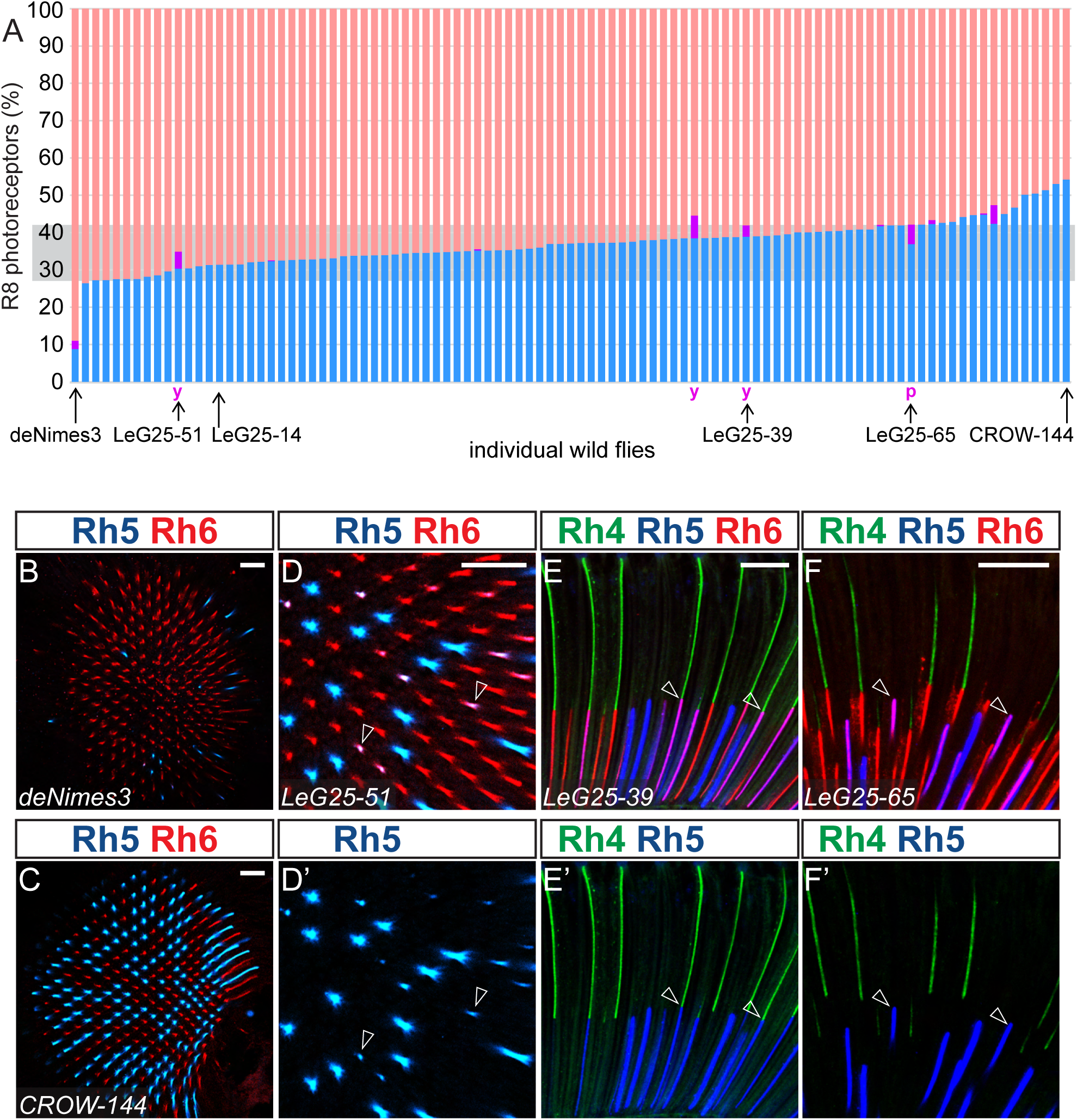
Distribution of Rh5/Rh6 phenotypes across wild flies. **A.** Quantified Rh5/Rh6 expression phenotypes of wild flies. Each vertical bar represents a single fly, with the fraction of Rh6-expressing R8s in red, Rh5-expressing R8s in blue, and Rh5/Rh6 co-expressing R8s in purple. Flies are arranged by increasing pR8 frequency. Rh5/Rh6 co-expressing R8s were assigned p or y identity whenever possible, based on Rh4 expression in the overlying R7 cells, as indicated by purple letters (see text for details). Flies shown in **B–F** are identified by name below the bars, as is LeG25-14, which is shown in Fig. 1B. The shaded band centred at ∼33% indicates the typical range of p:y ratios observed in laboratory wild-type strains. **B, C.** Wholemount retinas from wild flies illustrating the range of pR8:yR8 ratios, with the lowest ratio in deNimes3 (**B**) and the highest in CROW-144 (**C**). In CROW-144, pR8 photoreceptors are more abundant than yR8 photoreceptors. Compare with the wild-type retina in Fig. 1B. **D, D′.** Wholemount retina of LeG25-51 showing Rh5/Rh6 co-expression. Arrowheads mark co-expressing R8 photoreceptors. **D′** shows the same image with only the Rh5 channel. **E–F′.** Side views of ommatidia in wholemount retinas showing that Rh5/Rh6 co-expression can occur in yR8, as in LeG25-39 (**E, E′**), where Rh5/Rh6 co-expressing R8 photoreceptors contact Rh4-expressing yR7 photoreceptors (y-coexpression, e.g., arrowheads), or in pR8, as in LeG25-65 (**F, F′**), where they do not (p-coexpression, e.g., arrowheads). **E′** and **F′** show **E** and **F**, respectively, without the Rh6 channel. **B–D′** are single optical sections stained to reveal Rh5 (blue) and Rh6 (red). **E–F′** are Z-projections of 6 µm and 12 µm stacks, respectively, stained to reveal Rh4 (green), Rh5 (blue), and Rh6 (red). Scale bars, 20 µm.

Among the 97 flies examined, the majority showed the expected proportion of Rh5-expressing R8s (pR8), typically in the range of ∼30–40%, with no R8s co-expressing Rh5 and Rh6, corresponding to the typical laboratory wild-type phenotype. However, we also observed substantially different p:y R8 ratio phenotypes: in five flies, 50% or more of R8s expressed Rh5, while in one male fly, designated deNimes3, only ∼9% did (Fig. 2A–C). In six flies, co-expression of Rh5 and Rh6 occurred in 2–9% of R8s (Fig. 2A,D). In four flies, we were able to determine whether Rh5/Rh6 co-expressing R8s were paired with Rh4-expressing yR7s or with pR7s. In three of them, co-expression occurred in yR8s (y-coexpression), indicating ectopic Rh5 expression in yR8s (Fig. 2A,E), whereas in one fly, Rh5/Rh6 co-expression occurred in pR8s (p-coexpression), indicating ectopic Rh6 expression in these cells (Fig. 2A,F). These results show that Rh5/Rh6 expression patterns vary within wild populations, although it remains unclear whether this variation reflects genetic differences, environmental influences, or interactions between genetic and environmental factors.

### A loss-of-function sevenless allele underlies the deNimes3 phenotype

While ageing the flies, we also collected their progeny to establish isofemale or isomale lines. To test whether genetic variants contribute to the observed Rh5/Rh6 expression variation, we focused on the deNimes3 male fly. Among its progeny from mating with laboratory wild-type females, some flies showed a similarly low percentage of pR8s. Through a series of genetic crosses, we determined that the causative variant is recessive and X-linked and established a stable homozygous line carrying the mutation, which we called *deNimes* (Fig. 3A,B, Supplementary Data 1).

**Figure 3.**
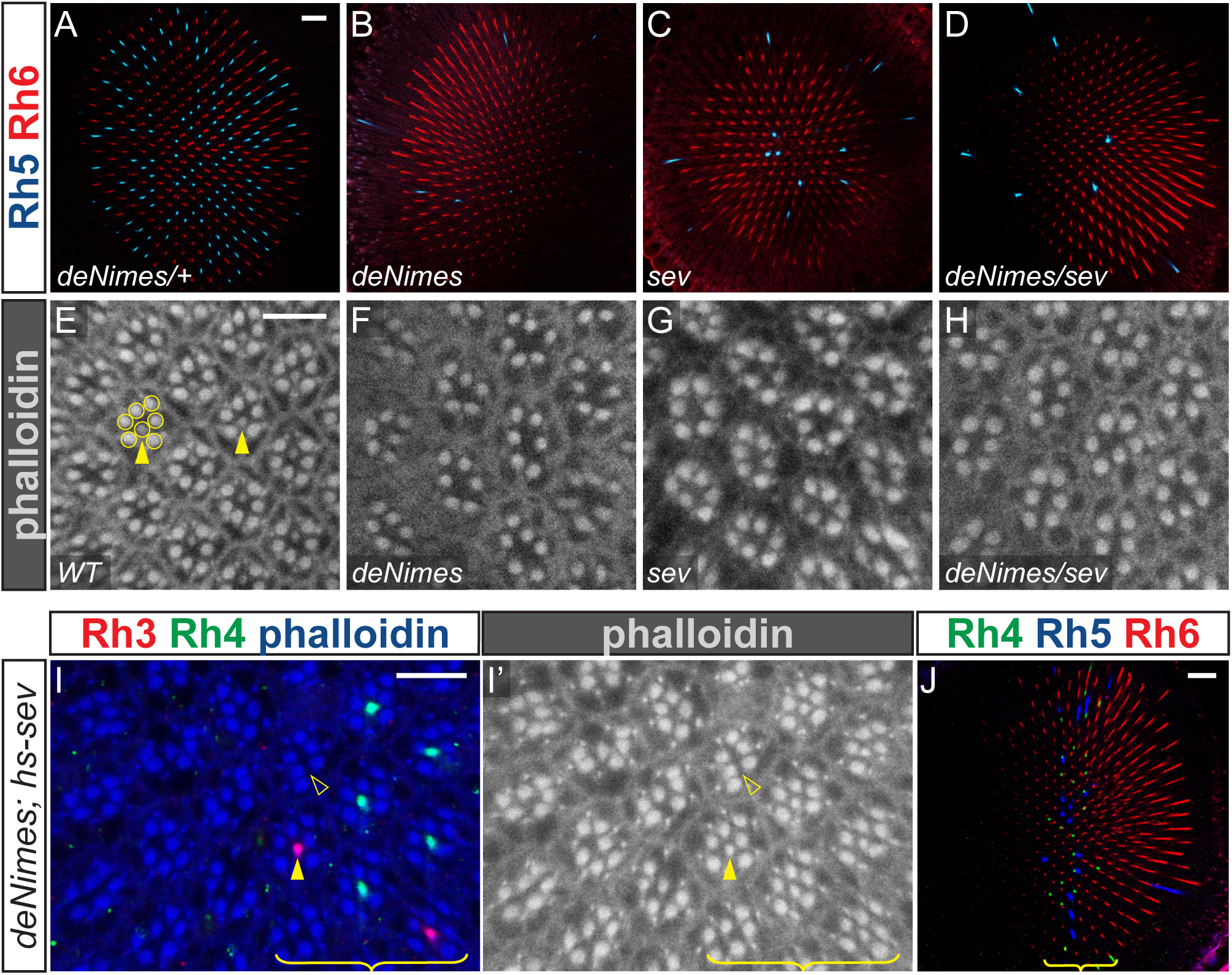
*deNimes* is a *sevenless* allele that eliminates R7 photoreceptors and is rescued by transient *sevenless* expression. A–D. Wholemount retinas stained to reveal Rh5 (blue) and Rh6 (red). *deNimes*/+ heterozygotes show a wild-type Rh5/Rh6 pattern (A; compare with Fig. 1B), whereas *deNimes* mutants show a reduced fraction of Rh5-expressing R8 photoreceptors (B). *sevenless* (*sev*) mutants show a similar Rh5/Rh6 phenotype (C), as do *deNimes*/*sev* trans-heterozygotes (D), indicating failure to complement. E–H. Optical sections at the R7 level stained with phalloidin to visualise rhabdomeres (photoreceptor light-sensing organelles). In wild-type retina (E), rhabdomeres of a single ommatidium are outlined. Filled arrowheads mark R7 rhabdomeres, identifiable by their central position and smaller diameter relative to R1–R6. R7 rhabdomeres are absent in *deNimes* (F) and *sev* (G). R7 rhabdomeres are also absent in *deNimes*/*sev* trans-heterozygotes (H), indicating failure to complement for the R7-loss phenotype. I, I′. Pulse expression of *sev* under the heat-shock promoter rescues R7 photoreceptors in *deNimes* mutants in a band spanning approximately four ommatidial columns. In I, Rh3 (red) and Rh4 (green) expression is shown together with phalloidin staining (blue). Six ommatidial columns are shown, with three columns lacking rescue on the left and three columns on the right (brace) containing many rescued ommatidia. An open arrowhead marks one ommatidium lacking R7, and a filled arrowhead marks one ommatidium with a rescued R7 expressing Rh3. I′ shows the same field with phalloidin staining only (white). J. Wholemount retina from a deNimes mutant rescued by the same pulse of heat-shock-induced *sev* expression as in I and I′, showing a vertical band (brace) of restored Rh4 (green) and Rh5 (blue) expression. Rh6-expressing cells are in red. The image is a composite of two superimposed optical sections acquired 22 µm apart at the R7 and R8 levels. Scale bars, 20 µm (A–D, J) and 10 µm (E–I′).

To assess possible effects of the variant on other Rhodopsins, we stained the retinas with antibodies against Rh3 and Rh4. Both were absent in *deNimes* flies, except for a row of Rh3-expressing R8 photoreceptors along the dorsal margin of the retina where both R7 and R8 normally express Rh3 (Supplementary Fig. 1A,B). Phalloidin staining confirmed that the mutant flies lacked R7 photoreceptors (Fig. 3E,F), similar to the loss-of-function of the *sevenless* (*sev*) gene^34,35^ (Fig. 3G, Supplementary Fig. 1C), which encodes a transmembrane receptor tyrosine kinase required to specify R7 photoreceptors^36,37^. Indeed, the *deNimes* mutation did not complement a *sev* null allele (Fig. 3D,H, Supplementary Fig. 1C). Furthermore, it was rescued by expression of a *sev* transgene in the same pulse-rescue paradigm used for *sev* mutants^37–39:^ because eye photoreceptors differentiate in a posterior-to-anterior wave, a 30-minute pulse of a heat-shock promoter-driven *sev* transgene during late larval stages rescued the loss of R7 in approximately four columns of ommatidia in the middle of the retina of *deNimes* mutants, restoring the presence of R7s with their expression of Rh3 and Rh4 (Fig. 3I), and also Rh5 expression in R8s (Fig. 3J).

In searching for the sequence change causing the *deNimes* phenotype, we identified an altered region within the *sev* locus. With four different primer pairs spanning this region, PCR products from *deNimes* flies appeared as smears rather than as the sharp bands seen in wild-type controls, with each smear corresponding in size to its wild-type counterpart while migrating slightly faster on the gel (Supplementary Fig. 1D). Sequencing the largest *deNimes* PCR product from either end yielded traces that terminated abruptly at specific positions (Supplementary Fig. 1E). In wild-type, these positions are separated by 72 bp. These results identify the mutation present in the original deNimes3 fly and its progeny as a sequence alteration replacing this short interval at the junction between the second exon (the first coding exon) and the following intron of *sev* (Supplementary Table 1).

### A screen for Rh5/Rh6 expression variants in DGRP2 lines

To determine the range of R8 photoreceptor differentiation and maintenance phenotypes that can arise from natural genetic variation in *Drosophila*, we examined Rh5 and Rh6 expression in the DGRP2 inbred fly lines. As for the wild flies described above, flies from the 205 available DGRP2 lines were aged for over two weeks and their retinas were stained for Rh5 and Rh6 expression. A phalloidin counterstain allowed us to identify R8 photoreceptors even when they were not expressing either Rh5 or Rh6.

We observed the same types of phenotypes as in the wild flies, but stronger, likely because these flies have been isogenised and are thus homozygous for the causative variants (Fig. 4A, Supplementary Data 1). We also observed complete loss of Rh5 or Rh6 phenotypes.

**Figure 4.**
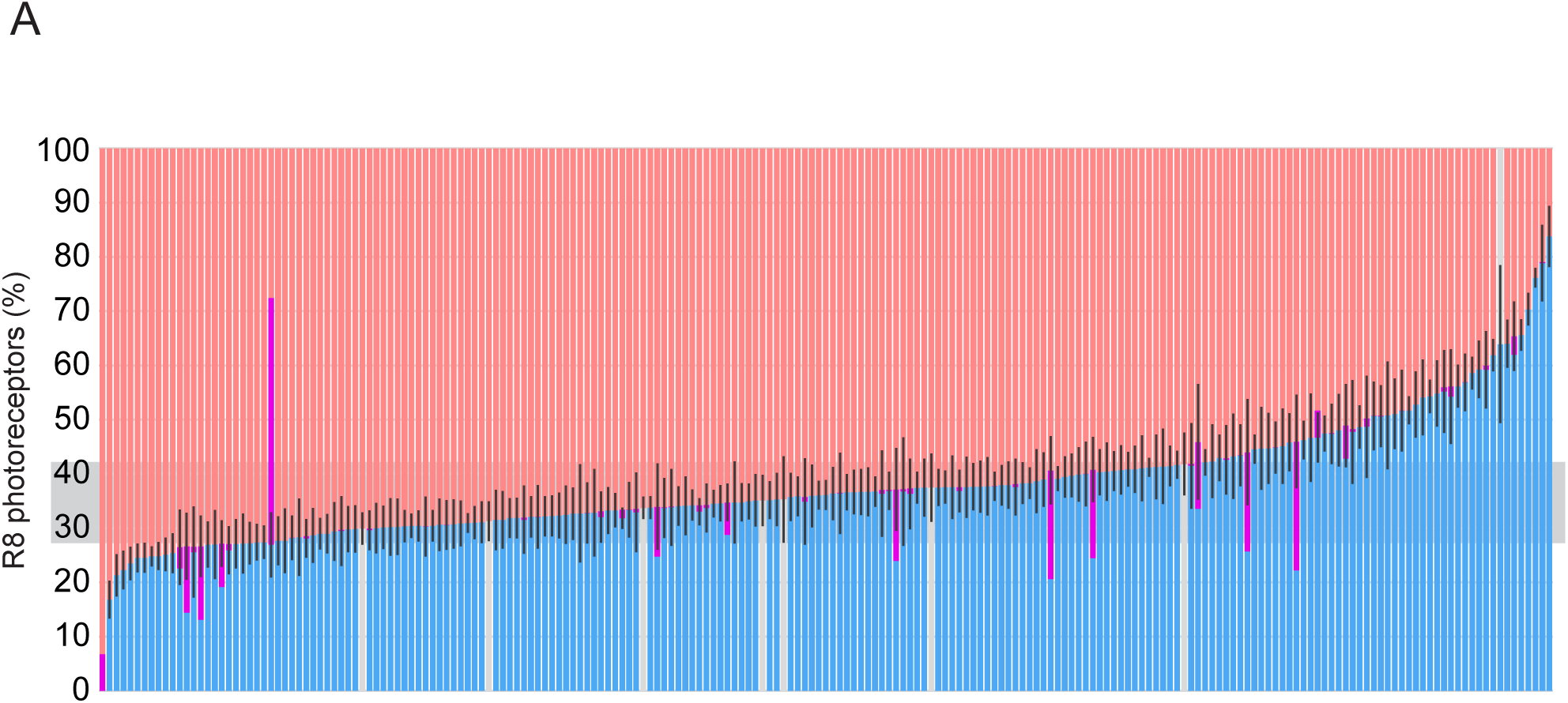
Distribution of Rh5/Rh6 phenotypes across the DGRP2 lines. A. Quantified Rh5/Rh6 expression phenotypes of the DGRP2 lines. Each vertical bar represents one DGRP line, with the fraction of Rh6-expressing R8s in red, Rh5-expressing R8s in blue, Rh5/Rh6 co-expressing R8s in purple, and R8s lacking detectable Rh5 or Rh6 in grey. Lines are ordered by increasing pR8 frequency (see Fig. 6 and associated text for details on classification of Rh5/Rh6 co-expressing R8s). Error bars indicate SD of pR8 frequency for each line. The shaded band centred at ∼33% indicates the typical range of p:y ratios observed in laboratory wild-type strains.

Strikingly, strong Rh5/Rh6 phenotypes were common, occurring in ∼1 in 6 DGRP lines. In the sections below, we systematically analyse Rh5/Rh6 phenotypes across the panel to classify them into the eight qualitative categories predicted in the Introduction and, where feasible, identify causal variants for selected examples.

### Loss of R8 Rhodopsin phenotypes: Lines with ‘empty’ R8 photoreceptors carry Rhodopsin mutations

Colour blindness due to loss or sequence alteration of an opsin gene is arguably the best-known human natural genetic variant class affecting the senses. Thus, we wanted to determine the causes of the loss of Rh5 or Rh6 expression that we observed in several lines (Fig. 4A).

Seven DGRP lines displayed a complete absence of detectable Rh5 protein but a normal number of photoreceptors that expressed Rh6 (Fig. 4A, Fig. 5A,D). The presence of a central R8 rhabdomere in these empty ommatidia indicated that this phenotype does not represent selective loss of pR8 photoreceptors. To identify the sequence change that causes the lack of Rh5 immunoreactivity, we first tested the possibility that *Rh5* itself was mutated. We crossed all seven lines with known *Rh5* null mutants that have the entire *Rh5* ORF removed^9^. The resulting trans-heterozygotes showed no Rh5 expression (Fig. 5B,C,E,F). Examination of the DGRP sequences of the *Rh5* locus and our own sequencing in the seven lines revealed that they carry two distinct *Rh5* coding variants. DGRP-397, DGRP-589 and DGRP-790 carry a C to T SNP (2L_12009649_SNP, here *Rh5^Q3^*^65^*\**, Supplementary Table 1) that converts the Q365 codon to a STOP codon. This truncates the C-terminal end of the Rh5 protein (Fig. 5G) and removes the epitope for the monoclonal anti-Rh5 antibody used in this study^40^. DGRP-559, DGRP-646, DGRP-832 and DGRP-900 carry a short 26 bp deletion (2L_12008602_DEL, here *Rh5^Δ^*^26^, Supplementary Table 1) in the first exon within the *Rh5* open reading frame (Fig. 5G), introducing a frameshift that results in termination of the ORF after 27 codons. *Rh5^Δ^*^26^ is likely a null allele of *Rh5*, as it removes most of the protein. These two *Rh5* alleles are not present in other DGRP lines, which all show at least some Rh5 expression. It is worth noting that the *Rh5^Δ26^* allele was correctly called by the DGRP computational pipeline only in the DGRP-832 line and not in the other three lines carrying it.

**Figure 5.**
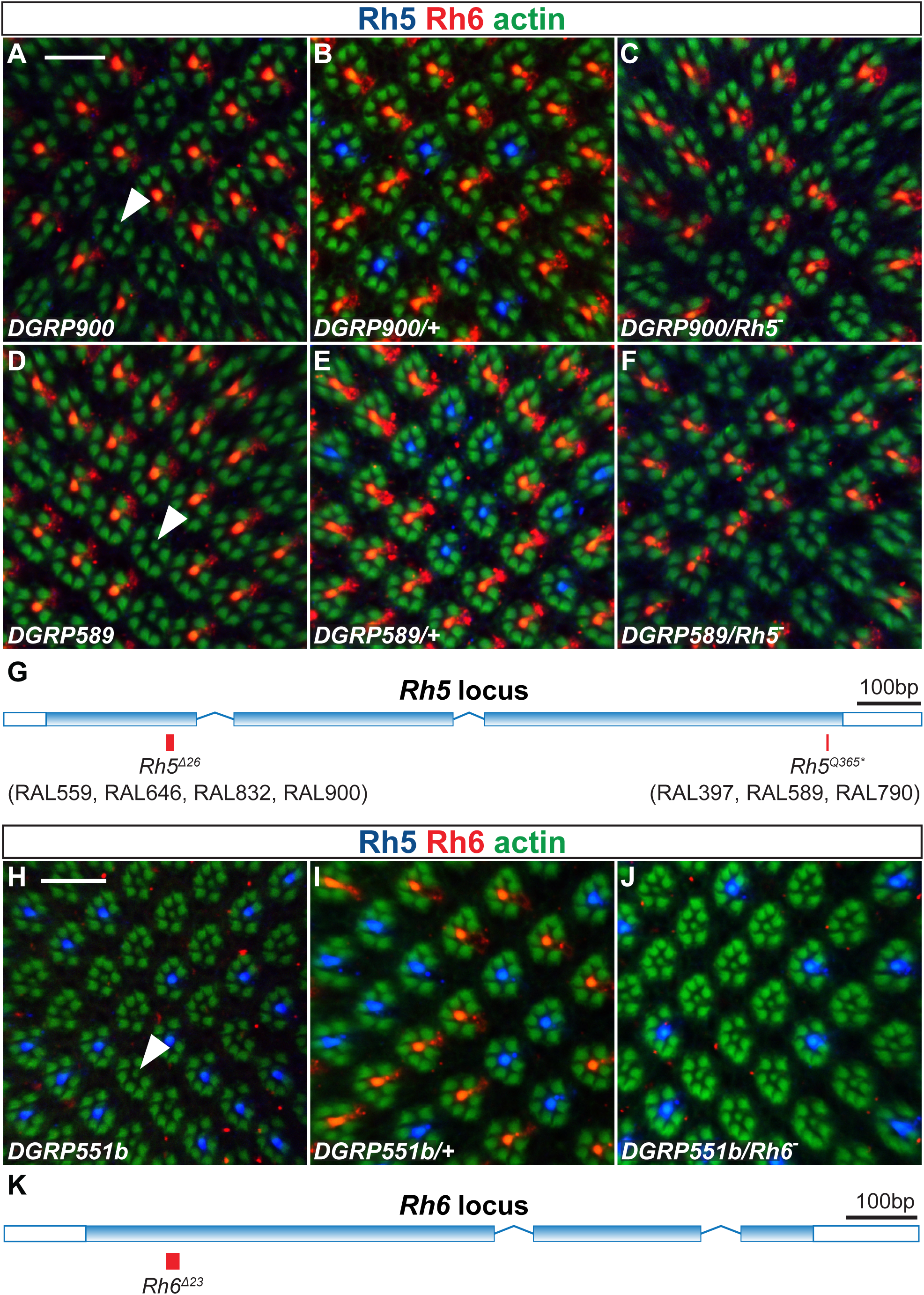
Absent Rh5 or Rh6 expression phenotypes among the DGRP lines are due to variants disrupting the open reading frames of these genes. A–C. DGRP-900 lacks Rh5 (blue) expression (A). The phenotype is recessive (heterozygotes, B) and fails to complement a *Rh5* null mutation (trans-heterozygotes, C). Presence of phalloidin-stained R8 rhabdomeres (e.g. arrowhead) indicates the presence of “empty” R8s. D–F. DGRP-589 lacks Rh5 expression (D). The phenotype is recessive (heterozygotes, E) and fails to complement a *Rh5* null mutation (trans-heterozygotes, F). Presence of phalloidin-stained R8 rhabdomeres (e.g. arrowhead) indicates the presence of “empty” R8s. G. Map of the Rh5 locus showing the position of the two *Rh5* ORF-disrupting variants, *Rh5^Δ26^* and *Rh5^Q365*^*. The DGRP lines carrying each variant are indicated. H–J. A variant abolishes Rh6 (red) expression in a line derived from DGRP-551 (H). The phenotype is recessive (heterozygotes, I) and fails to complement a Rh6 null mutation (trans-heterozygotes, J). Presence of phalloidin-stained R8 rhabdomeres (e.g. arrowhead) indicates the presence of “empty” R8s. K. Map of the *Rh6* locus showing the position of the Rh6 ORF-disrupting variant, *Rh6^Δ23^*. All images are optical sections through central retinas stained with anti-Rh5 (blue) and anti-Rh6 (red) antibodies and counterstained with phalloidin to visualise rhabdomeres (green). Scale bars, 10 µm.

We also observed the opposite phenotype: the DGRP-551 line consistently showed some, but not all, flies entirely without Rh6 expression^31^. We hypothesised that the mixture of phenotypes was due to genetic heterogeneity within this line. Thus, we set up several single-pair crosses and produced a sub-line in which all flies showed complete loss of Rh6 expression (Fig. 5H). Complementation experiments (Fig. 5I,J) and sequencing showed that the mutation was in the *Rh6* gene itself and identified a 23 bp deletion (here *Rh6^Δ23^*, Supplementary Table 1) within the *Rh6* open reading frame (Fig. 5K) that causes a frameshift after the Trp46 codon. By examining the original DGRP-551 sequencing reads^31,32^, we found that approximately a third of the reads covering this polymorphic locus showed the mutant sequence, indicating that this variant was present and segregating at the time it was sequenced soon after the line was established and is not a spontaneous mutation that arose more recently. Why both this allele and its wild-type counterpart have remained in the line since then remains a mystery.

Thus, genetic variants causing both Rh5 and Rh6 loss phenotypes are present among the DGRP lines and they directly affect the coding sequences of the two genes.

### Ectopic R8 Rhodopsin phenotypes: Natural variants causing Rh5/Rh6 co-expression

Normally, Rh5 and Rh6 are not co-expressed in the same R8; however, 26 DGRP lines showed different degrees (in 1%–45% of R8s) of Rh5/Rh6 co-expression (purple in Fig. 4A, Fig. 6C, Supplementary Data 1). This may represent two distinct phenotypes: co-expression may occur due to inappropriate expression of Rh6 in pR8 (p-coexpression) or due to inappropriate expression of Rh5 in yR8 (y-coexpression). To classify the DGRP co-expression phenotypes into these two categories, we tested whether Rh5/Rh6 co-expressing R8 photoreceptors were coupled with p or y R7 (Fig. 6A,B). Of the 16 lines with a fraction of co-expressing R8 greater than 3.5% (Fig. 6C), we found 9 lines in which co-expression occurred almost exclusively as p-coexpression (>98%), 2 lines in which co-expression occurred almost exclusively as y-coexpression (>98%), and 5 lines with some degree of both (Fig. 6D). In the latter group, co-expression could reflect either the presence of multiple variants or a single variant with mixed effects. These results show that genetic variants producing both classes of Rh5/Rh6 co-expression phenotypes, in pR8 and in yR8, are present among the DGRP lines.

**Figure 6.**
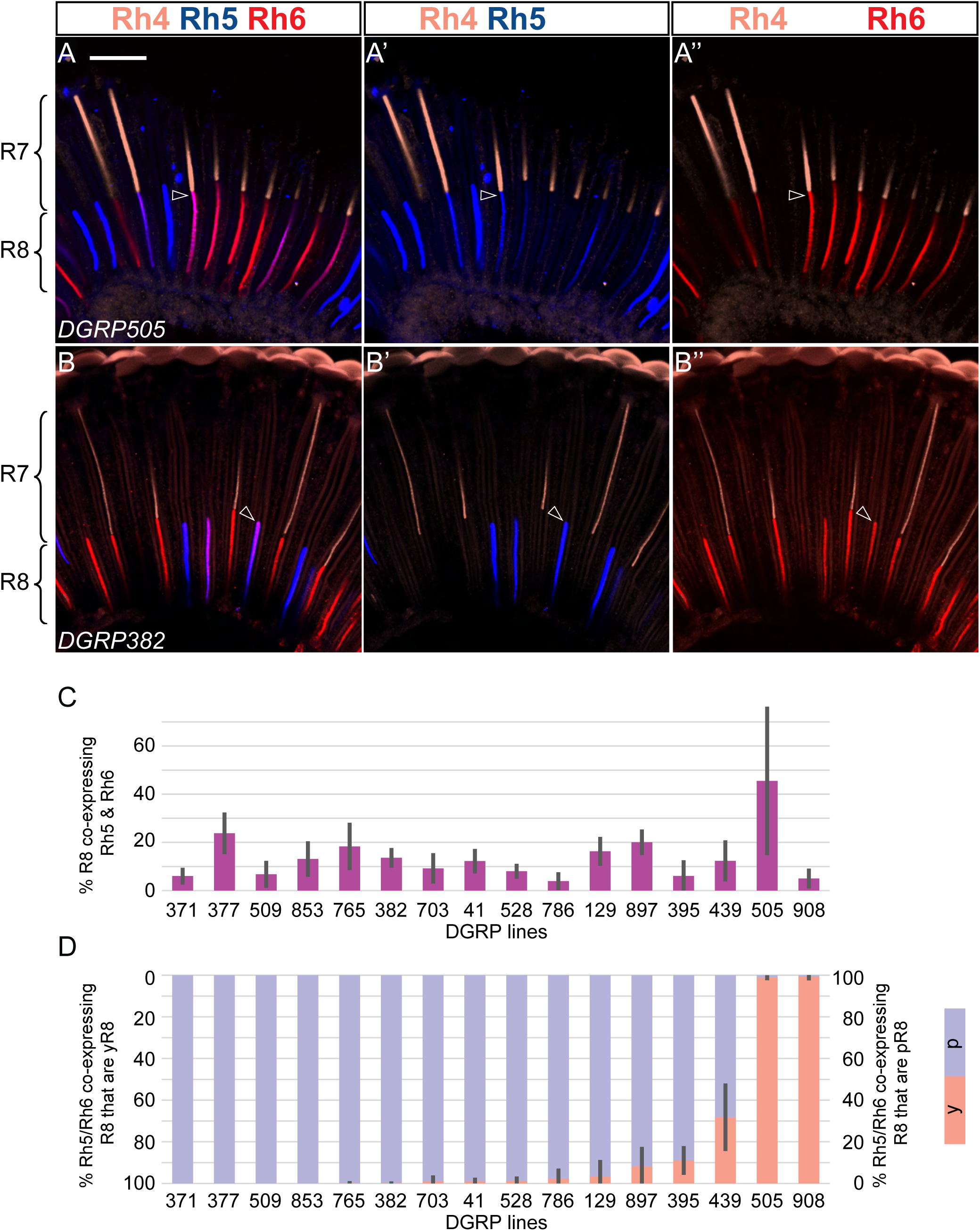
Rh5/Rh6 co-expression phenotypes among the DGRP lines. A. Side view of DGRP-505 ommatidia. Many R8 photoreceptors co-express Rh5 and Rh6 (e.g. arrowheads) and are paired with Rh4-expressing R7 photoreceptors. A′, A″, Image in (A) with the red and blue channels removed, respectively, to show Rh5 and Rh6 signal in the same R8 cells. B. Side view of DGRP-382 ommatidia. Many R8 photoreceptors co-express Rh5 and Rh6 (e.g. asterisk) and are paired with Rh4-negative R7 photoreceptors (R7 is not otherwise labelled). B′, B″, Image in (B) with the red and blue channels removed, respectively, to show Rh5 and Rh6 signal in the same R8 cells. C. Percentage of R8 photoreceptors co-expressing Rh5 and Rh6 among DGRP lines with co-expression in at least 3.5% of R8s. Error bars indicate SD. D. Proportion of Rh5/Rh6 co-expressing R8 photoreceptors that abut Rh4-expressing R7 photoreceptors (y-coexpression, red) or Rh4-negative R7 photoreceptors (p-coexpression, blue). All images are wholemount retinas stained with antibodies to reveal Rh4 (light red), Rh5 (blue), and Rh6 (red). (A), projection of a 4 µm stack, and (B), projection of a 5 µm stack of optical sections. Scale bar, 20 µm. Quantification in (D) was performed on z-stacks spanning R7–R8 contact points in the central retina and not on the illustrative images shown in (A) and (B).

### R8 subtype conversion phenotypes: Natural variants affecting the R8 p:y ratio

Given the conservation of the ∼1:2 p:y ommatidia ratio reported in Musca and Calliphora^18,41^, it was surprising to find substantial deviations from this ratio among the DGRP lines in both directions, ranging from 0% to 79% pR8 (Fig. 4A, Supplementary Data 1). Changes in the p:y R8 ratio, i.e. conversion phenotypes as defined in the Introduction, can occur either in coordination with or independently of changes in the R7 p:y ratio scored by Rh3 and Rh4 expression (Fig. 1D; see Introduction). Many conversion phenotypes among the DGRP lines are due to the *spineless* allele *sin* (in at least 47 lines) and other alleles of the ss transcription factor identified by Anderson et al.^16^ using a deficiency spanning *ss* to uncover the *ss* locus in each DGRP line; below, we will refer to these as the ss-screen variants. Because *ss* acts in R7 and is upstream of the p versus y subtype identity decision in R8 (Fig. 1C), these alleles are expected to cause coordinated conversion phenotypes. We therefore used the ss-screen to prioritise lines in our search for R7–R8 mis-coordination, focusing on those whose pR8:yR8 ratios could not be explained by ss-screen variants alone. We then compared Rhodopsin expression in R7 and R8 in these lines to distinguish coordinated from uncoordinated conversion phenotypes.

Specifically, we compared for each line the pR7:yR7 ratio obtained in the ss-screen with the pR8:yR8 ratio obtained in the present study (Fig. 7A, Supplementary Data 1). For most lines, the p:y ratios measured in the two screens were closely similar, confirming that ss-dependent shifts in R7 subtype identity are reflected quantitatively in R8 subtype ratios across natural genetic backgrounds. Lines with differing p:y ratios between the two screens were inferred to carry non-ss-screen variants affecting the pR8:yR8 ratio. Nine DGRP lines showed a significantly higher pR8:yR8 ratio than the pR7:yR7 ratio observed in the ss-screen (e.g. DGRP-437, DGRP-890), while four DGRP lines showed a significantly lower pR8:yR8 ratio (e.g. DGRP-509, DGRP-320).

**Figure 7.**
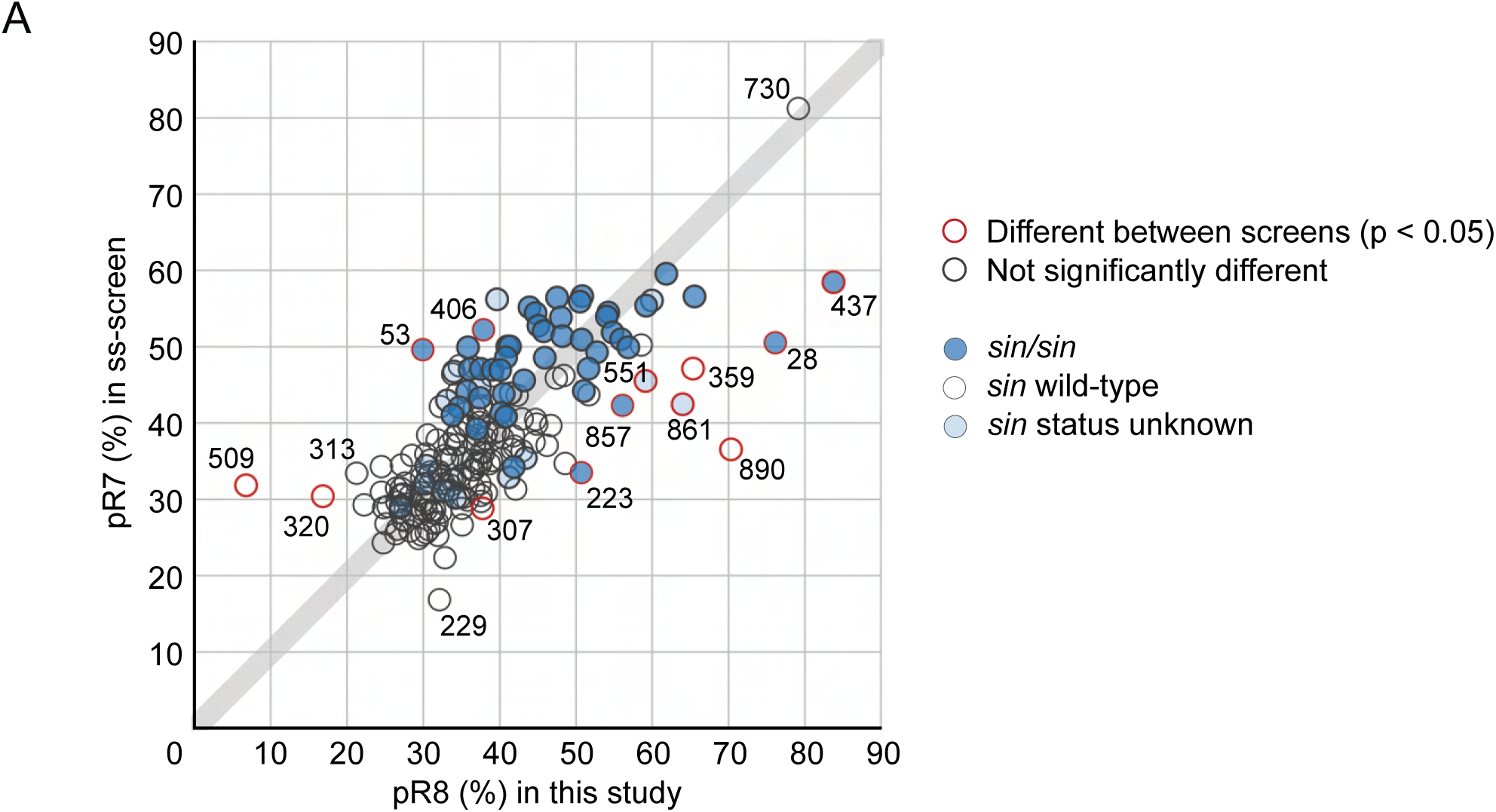
Comparison of R7 subtype ratios from the ss-screen and R8 subtype ratios from this study across the DGRP2 lines. A. For each DGRP line, %pR7 measured in the ss-screen (y axis) is plotted against %pR8 measured in this study (x axis). Divergence from the y = x line (grey) indicates differences between the two screens. Lines with significant differences are outlined in red. *sin* status is indicated by shading: lines with wild-type *sin* are white, homozygous *sin* lines are dark blue, and lines with undetermined *sin* status are light blue. Lines with significant differences or discussed in the text are labelled with DGRP numbers.

We next examined selected lines with differing p:y ratios in the two screens to determine whether they showed coordinated or mis-coordinated R7–R8 subtype identities. In lines with a higher pR8:yR8 ratio in our screen than the corresponding pR7:yR7 ratio in the ss-screen (below the y = x diagonal in Fig. 7A), R7–R8 mis-coordination would be expected to produce ommatidia containing Rh4-expressing R7s paired with Rh5-expressing R8s. We assessed R7–R8 coordination in eight such lines (Fig. 8A). All but one line showed coordinated R7–R8 subtype identities, with no Rh4–Rh5 mis-coordinated ommatidia, indicating that in these lines the p:y ratio shift reflects variants that affect R7 subtype identity but are not ss alleles. Only DGRP-890 exhibited Rh4–Rh5 ommatidia (Fig. 8B–E). The variant underlying this phenotype could affect pR7-to-R8 signalling or its consolidation in R8 (see Introduction) and is currently under investigation.

**Figure 8.**
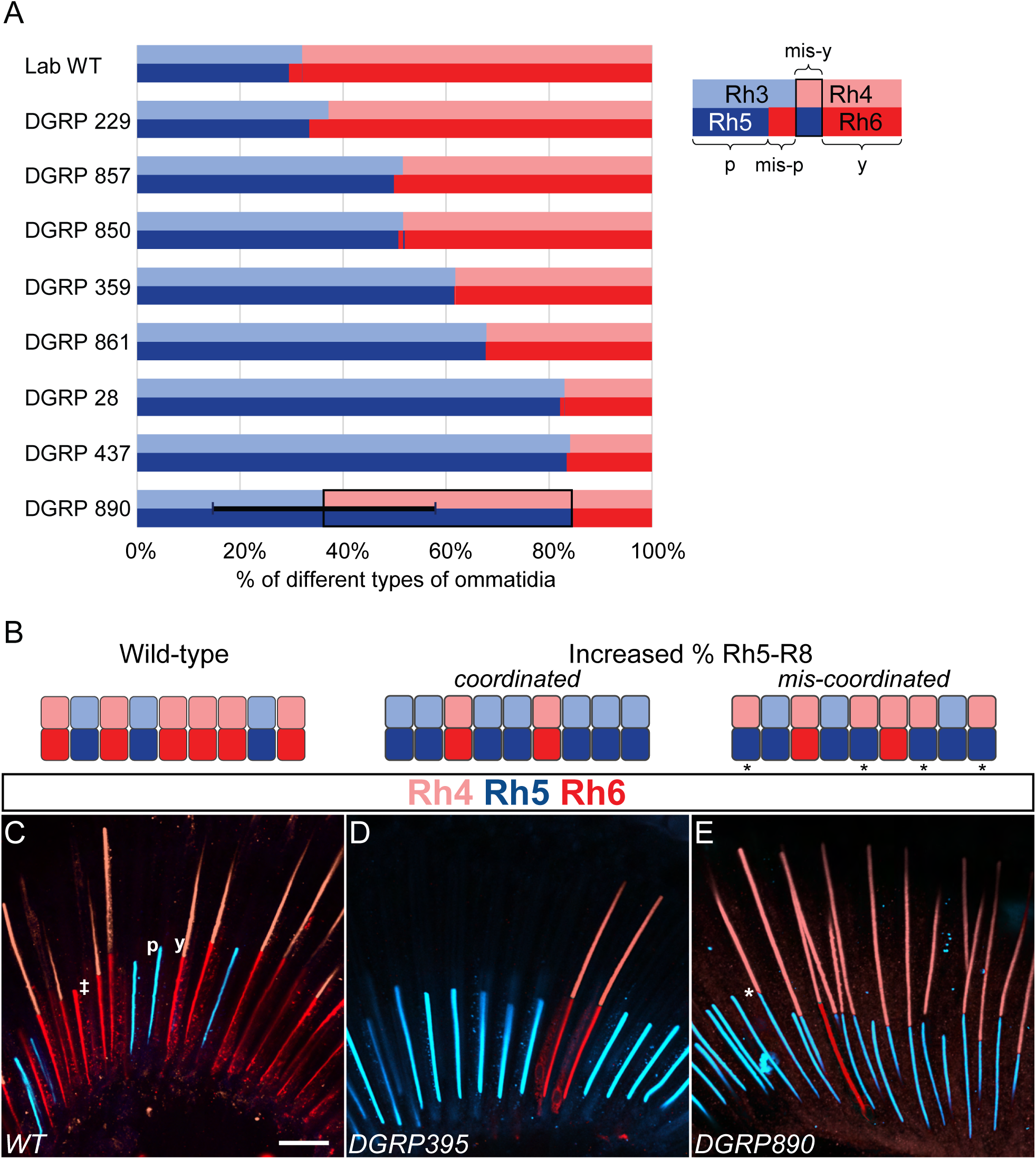
R7–R8 coordination and mis-y (Rh4–Rh5) ommatidia in selected DGRP lines. A. Percentages of normal and mis-coordinated ommatidia in selected DGRP lines with higher %pR8 in this study than %pR7 in the ss-screen. Ommatidia were scored from z-stacks stained for Rh4, Rh5, and Rh6. Rh3 expression was inferred for Rh4-negative R7. Each horizontal double-bar shows the fraction of four ommatidial types (p, mis-p, mis-y, y; total 100%), displayed as paired R7 (top) and R8 (bottom) Rhodopsin expression blocks (see in-panel key): p (Rh3–Rh5), y (Rh4–Rh6), mis-p (Rh3–Rh6), and mis-y (Rh4–Rh5). mis-y ommatidia were prominent only in DGRP-890 (outlined), and error bars indicate SD of %mis-y for DGRP-890. B. Schematics illustrating laboratory wild-type pattern and two phenotypes with an increased fraction of Rh5-expressing R8 photoreceptors, corresponding to the examples in C–E. In both phenotypes, not all ommatidia are shown as affected, to reflect the partial nature of these phenotypes in the corresponding lines. Increased Rh5 in R8 can be coordinated with an increased fraction of Rh3-expressing R7, preserving R7–R8 matching, or mis-coordinated, producing mis-y ommatidia (Rh4–Rh5). Colours as in A; asterisks mark mis-y ommatidia. C–E. Representative side views from lines quantified in (A), illustrating phenotypes with an increased fraction of Rh5-expressing R8 photoreceptors. In laboratory wild-type retinas (C), Rh4-expressing R7 are paired with Rh6-expressing R8 (y ommatidia), and occasional mis-p ommatidia (Rh3–Rh6) occur, here visible as Rh6-expressing R8 without a stained overlying R7 (marked with a double-dagger). In DGRP-359 (D), the increased fraction of Rh5-expressing R8 is accompanied by a corresponding increase in Rh3-expressing R7, and only correctly matched p (Rh3–Rh5) and y (Rh4–Rh6) ommatidia are observed. In contrast, in DGRP-890 (E), the increased fraction of Rh5-expressing R8 occurs without a corresponding increase in Rh3-expressing R7, producing mis-y ommatidia (Rh4–Rh5), one of which is marked with an asterisk. The micrographs are of wholemount retinas stained with antibodies to reveal expression of Rh4 in light red, Rh5 in blue, and Rh6 in red. (C), (D) and (E) are projections of 4 µm, 2 µm and 21 µm stacks of optical sections respectively. Scale bar, 20 µm.

Conversely, in lines with a lower pR8:yR8 ratio in our screen than the corresponding pR7:yR7 ratio in the ss-screen (above the y = x diagonal in Fig. 7A), R7–R8 mis-coordination would be expected to produce ommatidia containing Rh3-expressing R7s paired with Rh6-expressing R8s. Such mis-coordinated p-ommatidia occur at low frequency in wild-type laboratory flies, presumably reflecting occasional failures of pR7-to-R8 signalling to specify pR8 identity (Fig. 1C). Consistent with this, in our wild-type control line, 6.4% ± 3.0 SD of pR7s were paired with Rh6-expressing R8s (Fig. 9A).

**Figure 9.**
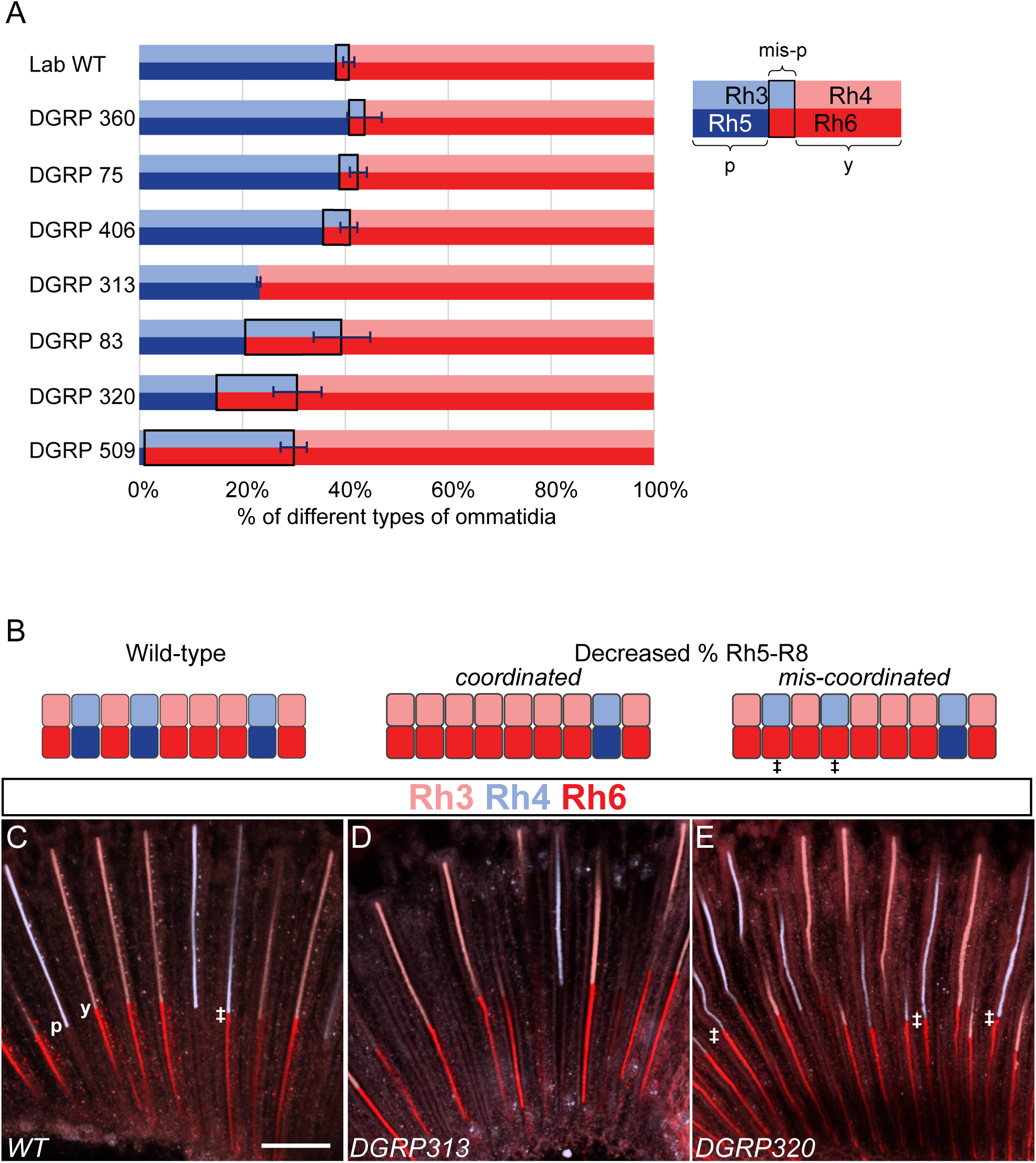
R7–R8 coordination and mis-p (Rh3–Rh6) ommatidia in selected DGRP lines. A. Percentages of normal and mis-coordinated ommatidia in selected DGRP lines with lower %pR8 in this study than %pR7 in the ss-screen. DGRP-313 was also included as an additional low-%pR8 line. Ommatidia were scored from z-stacks stained for Rh3, Rh4, and Rh6. Rh5 expression was inferred for Rh6-negative R8 photoreceptors. Each horizontal bar shows the fraction of three ommatidial types (p, mis-p, y; total 100%), displayed as paired R7 (top) and R8 (bottom) Rhodopsin expression blocks (see in-panel key): p (Rh3–Rh5), y (Rh4–Rh6), and mis-p (Rh3–Rh6), the only mis-coordinated type observed in these lines. Error bars indicate SD of %mis-p for each line. mis-p frequencies ranged from low in most lines to approximately half of Rh3-positive ommatidia in DGRP-83 and DGRP-320 and near-complete in DGRP-509. B. Schematics illustrating laboratory wild-type pattern and two phenotypes with a decreased fraction of Rh5-expressing R8 photoreceptors, corresponding to the examples in (C–E). In both phenotypes, not all ommatidia are shown as affected, to reflect the partial nature of these phenotypes in the corresponding lines. Decreased Rh5 in R8 can be coordinated with a decreased fraction of Rh3-expressing R7, preserving R7–R8 matching, or mis-coordinated, producing mis-p ommatidia (Rh3–Rh6). Colours as in A; double-daggers mark mis-p ommatidia. C–E. Representative side views from lines quantified in (A), illustrating phenotypes with a decreased fraction of Rh5-expressing R8 photoreceptors. In laboratory wild-type retinas (C), Rh4-expressing R7 are paired with Rh6-expressing R8 (y ommatidia), and occasional mis-p ommatidia (Rh3–Rh6) occur (marked with a double-dagger). In DGRP-313 (D), the decreased fraction of Rh5-expressing R8 is accompanied by a corresponding decrease in Rh3-expressing R7, and only correctly matched p (Rh3–Rh5) and y (Rh4–Rh6) ommatidia are observed. In contrast, in DGRP-320 (E), the decreased fraction of Rh5-expressing R8 occurs without a corresponding decrease in Rh3-expressing R7, producing frequent mis-p ommatidia (Rh3–Rh6) some of which are marked with double-daggers. All micrographs are of wholemount retinas stained with antibodies to reveal expression of Rh3 in light blue, Rh4 in light red, and Rh6 in red. (C), (D) and (E) are projections of 4 µm, 3 µm and 6 µm stacks of optical sections respectively. Scale bar, 20 µm.

We examined seven DGRP lines with pR8:yR8 ratios reduced relative to the ss-screen (Fig. 9A), four of which were significantly reduced (p < 0.05) (Fig. 7A). Across these lines, the frequency of mis-coordinated p-ommatidia, measured relative to pR7s, ranged from 0.6% to 97.2% of Rh3-expressing R7s paired with Rh6-expressing R8s. Three lines, DGRP-83, DGRP-320, and DGRP-509, showed particularly strong coordination defects, with 47.5% ± 12.9 SD, 51.1% ± 12.5 SD, and 97.2% ± 3.2 SD of pR7s paired with Rh6-expressing R8s, respectively (Fig. 9A–C,E). In these lines, the underlying variants strongly disrupt pR7-to-R8 signalling or its interpretation in R8. In the next section, we describe the identification of the variant responsible for the phenotype in DGRP-509.

As an example of a coordinated decrease in the p:y ratio, we examined the DGRP-313 line. It has 22.8% ± 4.8 SD pR8 (n = 21) and a negligible fraction of Rh3–Rh6 ommatidia (0.2% ± 0.3 SD) (Fig. 7A, Fig. 9A,B–D). For comparison, we selected two lines, DGRP-440 and DGRP-812, with pR8 frequencies close to the normal ∼30% (30.0% ± 4.7% SD, n = 14 and 30.6% ± 4.6% SD, n = 24, respectively) (Supplementary Data 1). The pR8:yR8 ratio of DGRP-313 is significantly lower than in either of these lines (p < 0.0001 and p < 0.000001, respectively; one-sided Welch’s t-tests). Thus, the DGRP2 collection contains at least one line carrying variant(s) associated with a coordinated reduction in the p:y ratio, affecting both R7 and R8 photoreceptor subtype frequencies.

In summary, DGRP lines carry genetic variants that alter the pR8:yR8 ratio in all four qualitative categories: increasing or decreasing it, each either in coordination with changes in R7 subtype identity or independently of R7s. In addition, coordinated increases in the pR8:yR8 ratio can arise from either *ss* alleles or variants affecting other genes that influence the pR7:yR7 ratio.

### A 214 bp deletion in the melt gene causes R8 subtype conversion in DGRP-509

To identify the genetic variant causing the DGRP-509 phenotype, we looked for mutations in candidate genes. The retinas of DGRP-509 have almost no Rh5-expressing photoreceptors and any weak Rh5 is co-expressed with Rh6 (Fig. 10B), while the Rh3/Rh4 expression appears normal (Fig. 9A, Fig. 10A,B). These observations indicate that the DGRP-509 phenotype reflects disruption of steps downstream of the p vs y fate decision by R7. These steps could, for example, be part of the R7-to-R8 signal^24^, or part of the Hippo/Melted bistable p vs y R8 switch^25–27^ (Fig. 1C). Loss of function of the *melted* (*melt*) gene^42^, a repressor of the Hippo pathway in the R8 photoreceptors, leads to a phenotype remarkably similar to that of the DGRP-509 line^27^ (Fig. 10C). Indeed, the mutation causing the DGRP-509 phenotype failed to complement a *melt* null mutation (Fig. 10D,E).

**Figure 10.**
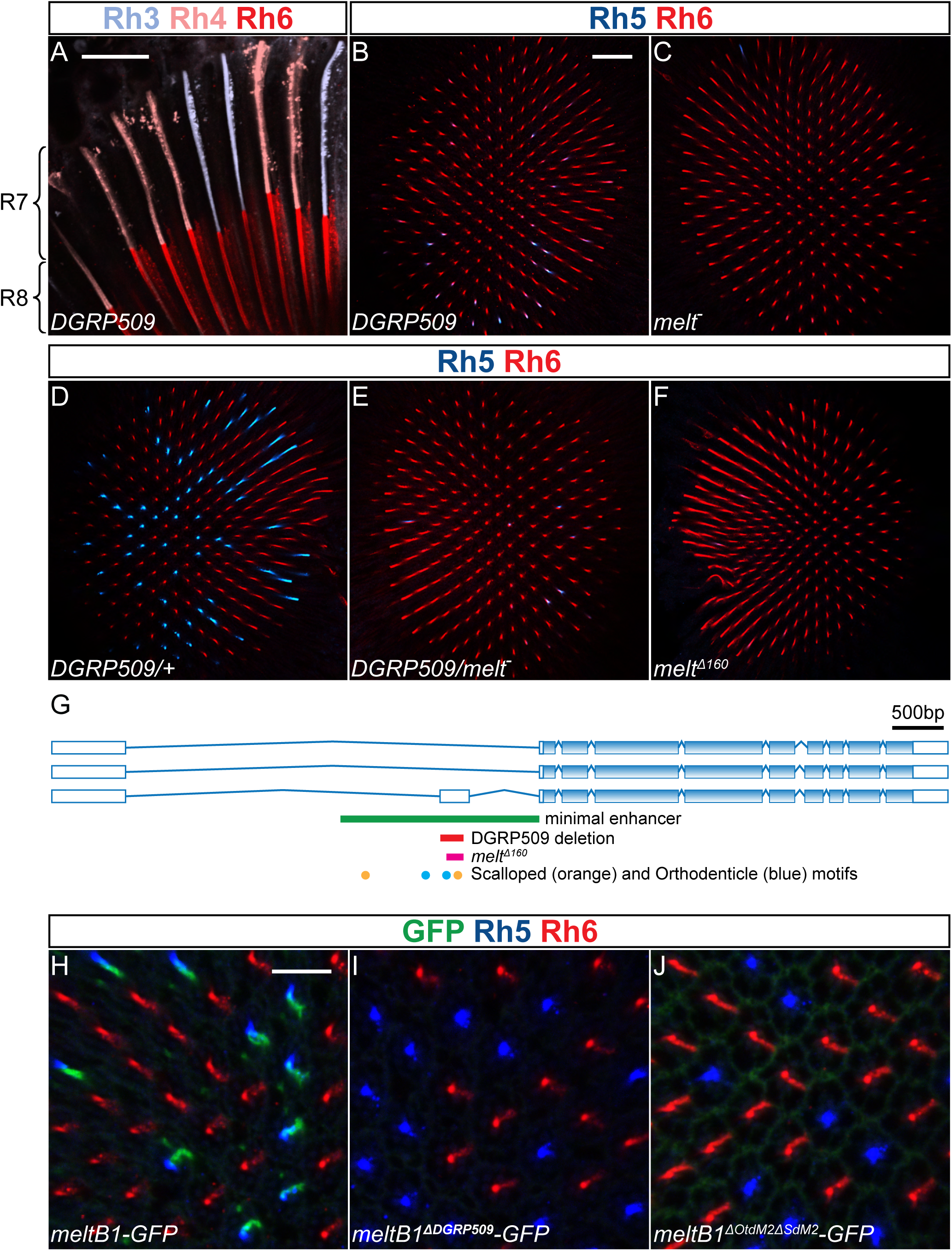
DGRP-509 phenotype is due to a deletion within the *melt* locus. A. DGRP-509 ommatidia, side view. Nearly all Rh3-expressing R7 (light blue) are paired with Rh6-expressing R8 (red), producing mis-p ommatidia (Rh3–Rh6). Rh4-expressing R7 (light red) paired with Rh6-expressing R8 represent normal y ommatidia (Rh4–Rh6). Compare with wild-type control in Fig. 9C. B. Nearly all R8 in DGRP-509 express Rh6. Most Rh5 expression is as co-expression with Rh6. C. Homozygous melt mutants have a similar phenotype to DGRP-509. D–E. The variant causing the DGRP-509 phenotype is recessive (D) and fails to complement a *melt* mutation (E). F. A 160 bp deletion within a *melt* intron (see (G) and text) phenocopies the variant causing the DGRP-509 phenotype. G. The *melt* locus. Three predicted alternatively spliced melt transcripts are shown, with coding sequence in blue. DGRP-509 carries a 214 bp deletion (red bar) within the large intron of melt. This sequence lies within the melt enhancer (green bar), sufficient to drive a GFP reporter in pR8. The CRISPR-generated 160 bp deletion (purple bar) is shorter at the 5′ end and matches the 3′ end of the DGRP-509 deletion. Both deletions disrupt one predicted Scalloped binding motif (orange dot) and one predicted Orthodenticle binding motif (blue dot) contained within the melt enhancer. H. The melt enhancer drives transgenic GFP reporter expression (green) in pR8. I. This expression is disrupted by the 214 bp DGRP-509 deletion within the transgene. J. This expression is also disrupted by a double mutation within the transgene of one Scalloped and one Orthodenticle motif contained in the 214 bp DGRP-509 deletion. All micrographs are of wholemount retinas stained with antibodies to reveal expression of Rh3 in light blue, Rh4 in light red, Rh5 in blue, Rh6 in red, and GFP in green, as indicated above each panel. (A), projection of a 2 µm stack of optical sections. Scale bar, 20 µm. (B– F), optical sections through central retinas. Scale bar, 25 µm. (H–J), zoomed-in optical sections within the central retina. Scale bar, 10 µm.

We examined annotated sequence variants in the *melt* locus reported by the DGRP consortium^31,32^, but none of them were unique to the DGRP-509 line. Nevertheless, we more closely examined the *melt* DGRP sequencing reads of the DGRP-509 line and, together with our own sequencing, we identified a 214 bp deletion within the first large intron of *melt* that was not present in any other DGRP line (Fig. 10G). This deletion is within the 2043 bp region comprising a *melt* enhancer (Fig. 10G) that is sufficient to drive transgenic GFP reporter (*meltB1*-GFP) expression in pR8 photoreceptors^25^ (Fig. 10H).

To determine whether the 214 bp deletion in DGRP-509 was responsible for the loss of Rh5 and gain of Rh6, we used CRISPR-Cas9 to create a deletion in the same region in an independent wild-type stock. We obtained a smaller 160 bp deletion, which left the first 54 bp of the 214 bp sequence intact. Nevertheless, this shorter deletion also caused the *melt* phenotype (Fig. 10F), indicating that the 160 bp region is necessary for *melt* expression and pR8 fate. As expected, this deletion also failed to complement the DGRP-509 mutation and the null *melt* allele (Supplementary Fig. 2A–C). Thus, the 214 bp deletion is the naturally occurring variant causing the abnormal R8 photoreceptor differentiation within the DGRP-509 line (Supplementary Table 1).

We also introduced the mutation in the *melt* enhancer reporter, *meltB1*-GFP that is expressed in pR8 photoreceptors^25^ (Fig. 10G–I). Consistent with DGRP-509, the 214 bp deletion also abolished the expression of this *melt* reporter in a wild-type background, confirming the essential role that this *cis-*regulatory region plays in *melt* activation. Thus, the DGRP-509 line allowed us to define, within the context of the endogenous *melt* locus, a critical enhancer region that is required for the correct Rh5/Rh6 expression pattern. Next, we sought to identify within this region the *cis*-regulatory motifs required for *melt* expression in the R8 photoreceptors. Two transcription factors, Scalloped (Sd) and Orthodenticle (Otd), are required for *melt* expression and the Rh5 fate^25^. Indeed, the 214 bp sequence deleted in *melt* in the DGRP-509 line contains one predicted Sd binding motif and one predicted K_50_/Otd binding motif^43^, and both of these motifs were also disrupted by the CRISPR-Cas9-generated 160 bp deletion. Specifically mutating these motifs also abolished reporter expression (Fig. 10J). Thus, the Rh5/Rh6 expression phenotype observed in the DGRP-509 line likely results from the loss of *melt* expression due to the removal by the 214 bp deletion of a pair of motifs, one Otd and one Sd, which are necessary for the pR8-specific *melt* expression.

## DISCUSSION

The well-characterised R7–R8 differentiation program produces a stereotyped Rhodopsin expression pattern: R8 photoreceptors normally express either Rh5 or Rh6, with their choice coordinated with the identity of the overlying R7. These simple rules allowed us to define the qualitative phenotypic space of Rh5/Rh6 expression as eight predicted outcomes. We used this framework to assess the scope of endophenotypic variation in genotypes derived from natural populations. We also identified several causal natural variants acting at different stages of photoreceptor differentiation.

We observed substantial variation in Rh5/Rh6 expression patterns, both in flies caught or conceived in the wild and across the DGRP2 panel, a snapshot of standing genetic variation from a single natural population in Raleigh, North Carolina, sampled in 2003. Remarkably, all eight predicted qualitative Rh5/Rh6 expression phenotypes were represented among the 205 DGRP2 lines. One of these, Rh5/Rh6 co-expression in pR8s, had not previously been reported in laboratory-generated mutants. Its occurrence among naturally derived genotypes illustrates how surveys of natural variation can extend the range of phenotypes described, even for a well-characterised developmental system. Phenotypes in the DGRP2 were generally more pronounced than in wild flies, possibly in part because of homozygosity; approximately one in six lines showed a strong phenotype. However, strong phenotypes were also present in wild flies, reflecting natural, outbred conditions. For example, the extreme p:y R8 ratio, which we mapped to a loss-of-function allele of *sevenless*, was first observed in the retina of an adult male caught in the wild (deNimes3). Together, our two phenotypic datasets demonstrate significant heritable variation in Rh5/Rh6 expression in natural *Drosophila melanogaster* populations.

In many of the phenotypes we examined, only a fraction of R8 photoreceptors are affected. As a result, categorical Rh5/Rh6 expression states at the single-cell level translate into graded, quantitative phenotypes at the level of the whole eye, which can be measured precisely and vary continuously in strength across genetic backgrounds. This feature is shared by many laboratory-generated mutations affecting Rh5/Rh6 expression, including null alleles^20–30^. For example, despite the complete absence of R7 photoreceptors in *sevenless* mutants, Rh5 expression persists in a subset of R8 cells for reasons that remain unclear.

This illustrates how a severe disruption at one developmental level can manifest as a partial, quantitative phenotype at another. The Rh5/Rh6 endophenotype thus captures an intermediate step between genotype and its consequences for visual function and, ultimately, fitness.

The causal variants we identified include both coding changes and regulatory mutations acting at different levels of the R7–R8 differentiation program. In the wild-caught deNimes3 fly, we identified a loss-of-function allele of *sevenless* that prevents R7 specification and thereby alters R8 differentiation. We traced the sequence alteration to a region spanning the exon–intron junction of the first coding exon. In the DGRP2 lines, we found loss-of-function frameshifts in *Rh5* and *Rh6*, and a second *Rh5* allele that truncates the C-terminal tail, removing the epitope recognised by our antibody, although its effect on Rh5 function remains uncertain. Particularly informative was a 214 bp deletion in an intronic enhancer of *melted* that severely disrupts pR8 differentiation without altering R7 identity. Together with the previously described regulatory *spineless* allele *sin*, these examples show that variants in genes known from laboratory studies do contribute to natural variation in Rh5/Rh6 expression. More broadly, they illustrate the value of studying a well-characterised endophenotype for identifying causal variants underlying natural phenotypic differences, a task that remains challenging even in *Drosophila*.

Even when the underlying genetic variants remain unidentified, the relationship between R7 and R8 identities can help locate their effects within the differentiation cascade. Changes in the p:y R8 ratio accompanied by corresponding changes in the p:y R7 ratio are consistent with perturbations of the stochastic *spineless* (*ss*) decision in R7. The previously described *ss* allele *sin*^16^ accounts for a substantial fraction of the lines with increased p:y R8 ratios, while other coordinated shifts could reflect additional *ss* alleles or variation in regulators of *ss* expression or function. In contrast, uncoordinated changes in the p:y R8 ratio, without corresponding changes in R7 identity, point to perturbations downstream of the R7 fate decision, either in the signal from pR7 to R8 or in its interpretation within R8, as illustrated by the *melted* enhancer deletion.

Loss of Rh5 or Rh6 expression that leaves R8 photoreceptors devoid of detectable Rhodopsin could result from mutations in the Rhodopsin genes themselves, as observed here, or from failures in specification, maintenance, or protein expression. Rh5/Rh6 co-expression, on the other hand, indicates disruption of the mechanisms that normally enforce mutually exclusive Rh5/Rh6 expression in R8 photoreceptors. Co-expression phenotypes could arise from defects in fate stabilisation or long-term maintenance, but they may also reflect alterations at the level of specification, for example if the output of the Rh5/Rh6 bistable switch is selectively compromised for one Rhodopsin. Interestingly, prolonged darkness can induce Rh5 expression in Rh6-expressing R8s^30^, showing that environmental conditions can also alter this expression pattern. Distinguishing whether the co-expression phenotypes observed here arise during photoreceptor specification or subsequent maintenance will require examining Rhodopsin expression in young flies.

We often observed combinations of Rh5/Rh6 phenotypes within the same line. For example, Rh5/Rh6 co-expression and loss of Rh5 or Rh6 staining were observed across a wide range of p:y R8 ratios, and p- and y-coexpression could occur together. The *ss* allele *sin* appears to interact with other variants that shift the p:y R8 ratio. Thus, the eight qualitative Rh5/Rh6 phenotypes described here can combine within the same genetic background to produce more complex whole-retina phenotypes. They could also combine with other photoreceptor phenotypes not examined here, such as variation in the expression of other Rhodopsins, photoreceptor morphology, or neuronal connectivity, either across the whole retina or in spatially restricted regions.

Several types of Rhodopsin expression changes that we observed occur as evolved retinal patterns across fly species. In the dorsal half of the *Drosophila* retina, yR7 photoreceptors co-express Rh3 and Rh4^44^, showing that mutually exclusive Rhodopsin expression can be relaxed in a spatially restricted manner. In the Dorsal Rim Area (DRA), both R7 and R8 express Rh3, while Rh5 and Rh6 are not expressed in R8^45,46^. Together with accompanying cellular specialisations in R7 and R8, this Rhodopsin expression pattern supports DRA-mediated polarised light vision. Hilbrant et al. identified a *D. simulans* strain with a dorsal-posterior patch enriched for pR7-containing ommatidia, illustrating natural variation in the spatial distribution of ommatidial subtypes^14^. In the housefly *Musca domestica*, *Rh4* has been lost during evolution^18,47^, and yR7 photoreceptors instead express *Rh2*^19^, which in *Drosophila* is expressed only in the ocelli. These examples illustrate how changes resembling the phenotypes uncovered here—including Rhodopsin co-expression, loss and replacement, and shifts in ommatidial subtype distribution—have contributed to the evolution of retinal organisation, often in combination with regional and cellular specialisations.

Developmental systems must be simultaneously robust enough to support normal organismal development and labile enough to evolve^48^. It is possible that the endophenotypic variation we observed reflects tolerance for deviation from a developmental optimum, which may have facilitated the evolution of divergent retinal phenotypes in other contexts. However, we do not know the extent to which the phenotypic differences we observed affect vision and ultimately fitness, even as we demonstrate that the R8 endophenotype is not robust to specific mutations^49^. The five causal variants we identified are all likely loss-of-function alleles and occurred at low frequencies in our samples, consistent with a trait under stabilising selection and the expectation of rare deleterious alleles maintained by mutation–selection balance for morphological traits^50^. This contrasts with the maintenance of genetic variation by balancing selection, which may dominate *Drosophila melanogaster* populations for life history traits^51–54^. We suggest that natural populations harbour alleles of moderate to large effect on R8 photoreceptor endophenotypes, representing deviations from a developmental optimum, with unknown but likely attenuated effects on functional vision.

The Rh5/Rh6 expression phenotypes described here have interesting parallels in the human retina. Although fly and human eyes differ greatly in their organisation, many genes involved in eye development are conserved, and a large proportion of genes implicated in human retinal diseases have *Drosophila* orthologs^55,56^. Variation in the p:y R8 ratio has a loose counterpart in the human retina, where the ratio of long-wavelength-sensitive (L) to medium-wavelength-sensitive (M) cones has been reported to vary by as much as 66-fold among individuals^57^. Genetic variants at the L/M-opsin and *NR2F2* loci have been associated with differences in these ratios^58,59^. Similarly, the *Rh5* and *Rh6* coding alleles identified here parallel human opsin mutations underlying common forms of colour vision deficiency^60^.

Finally, inherited retinal diseases provide many examples of individually rare alleles with strong phenotypic effects^61,62^, paralleling the pronounced Rh5/Rh6 phenotypes caused by some of the mutations identified here. Beyond retinal patterning, *Drosophila* also exhibits substantial heritable variation in phototaxis, a measure of visual behaviour. Of the 49 functionally validated candidate genes associated with this trait, 40 have human orthologs, seven of which are implicated in vision disorders^63^. Together, these parallels support the use of *Drosophila* as a model for investigating the genetic basis of natural variation in visual traits across species, including humans.

## MATERIALS AND METHODS

### Fly husbandry

Flies were raised and aged on standard cornmeal/molasses/agar medium at 24°C in ambient laboratory light unless otherwise specified.

### Antibodies

Antibodies and dilutions used were as follows: mouse anti-Rh3 (1:10) and mouse anti-Rh5 (1:100) (gifts from S. Britt); rat anti-Rh4 (1:100) (gift from C. Desplan); rabbit anti-Rh6 (1:2,000) (Ref. ^64^); sheep anti-GFP (1:500) (AbD Serotec). Secondary antibodies (donkey and goat) were Alexa Fluor-conjugated (Alexa Fluor 488, 1:1,000; Alexa Fluor 555, 1:750; Alexa Fluor 647, 1:500; Molecular Probes/Thermo Fisher). Alexa Fluor 488-conjugated or Alexa Fluor 647-conjugated phalloidin was used to visualise rhabdomeres (1:100; Molecular Probes/Thermo Fisher).

### Immunostaining

Immunostaining was performed as described in Ref. ^65^. Briefly, adult retinas were dissected out in phosphate-buffered saline (PBS), fixed for 15 min with 4% formaldehyde at room temperature, washed twice in PBS, and incubated with the primary antibodies diluted in blocking solution (PBS, 0.1% Triton X-100, 2% horse serum) for 2 days at 4°C. After two rinses and two 1 h washes with PBT (PBS, 0.3% Triton X-100), the retinas were incubated for 2 days at 4°C with secondary antibodies and phalloidin diluted in blocking solution.

Retinas were rinsed twice and, after two 1 h washes with PBT, mounted in SlowFade Gold (Invitrogen/Thermo Fisher).

### Confocal imaging and image processing

Samples were imaged using Leica TCS SP5 and SP8 confocal microscopes. For cross-sectional views of photoreceptors, optical sections were acquired approximately 10 µm distal to R8 nuclei in the central retina. For side views of photoreceptors, a short stack of optical sections was acquired near the rim of the retina and shown as a z-projection. Note that side-view images were used for illustrations only and were not used for quantification. For comparison of R8 and R7 Rhodopsin expression, a stack capturing both R8 and R7 layers was acquired. Images were processed using Leica Confocal Software (LCS), Adobe Photoshop, and Fiji software.

### Quantification

Rh5-expressing, Rh6-expressing, Rh5/Rh6-coexpressing, and “empty” R8s were counted in Fiji using the Cell Counter plugin. “Empty” R8s (no Rh5 or Rh6 staining) were defined as ommatidia (visualised by phalloidin, e.g. see Fig. 5A) within the Rh5-positive region of the retina in Rh6 mutants or within the Rh6-positive region in Rh5 mutants. For R7–R8 comparisons, only ommatidia with both R7 and R8 visible in the same z-stack were counted. Ommatidia were classified by Rhodopsin expression in R7 and R8. In some analyses, only specific ommatidial classes were counted (e.g., to determine the percentage of Rh5/Rh6-coexpressing R8s that were in p or y ommatidia, only ommatidia containing a Rh5/Rh6-coexpressing R8 were scored). Each fly was quantified once. For most experiments, one retina was analysed per fly. For wild flies, counts from both retinas, when available, were pooled to obtain a single measurement per individual. In all cases, individual flies were treated as independent biological replicates. The number of individuals per genotype and the number of R8 photoreceptors per retina are provided in the raw count datasets (Supplementary Data 1).

### Wild flies

Wild flies were obtained from three locations. Flies from Saint-Bauzély, France, were caught in August and November 2017. Flies from Le Gouray, France, were caught in August 2020 and July 2025. In both cases the flies were aged for at least two weeks after capture (at room temperature, near a window under natural daylight) before phenotyping. These are referred to as wild-caught flies. Flies from Crowborough, UK, were caught in July 2025. Because many of them showed Rh5/Rh6 co-expression, potentially due to prolonged shipment in darkness^30^, we instead phenotyped one 2-week-old daughter from each wild-mated female; these are referred to as wild-conceived flies. Wild-caught and wild-conceived are referred to collectively as wild flies. All wild flies were stained with antibodies to visualise Rh5 and Rh6 and, in some experiments, Rh4. See Supplementary Data 1 for details on each fly and quantitation of the phenotypes.

### deNimes3 male fly and its progeny

The wild-caught “deNimes3” male fly from the Saint-Bauzély collection was crossed to yw; If/CyO; TM2/TM6b balancer stock. The reduced p:y R8 ratio phenotype was not observed in the F1 progeny but was observed in some F2 progeny. Based on subsequent crosses, the mutation (*deNimes)* was inferred to be X-linked recessive. A stable line homozygous for the mutant X chromosome was established by crossing phenotypically verified *deNimes* males to females carrying the FM7 balancer X chromosome. For *sevenless* (*sev*) complementation experiments, *deNimes/sev*^14^ females were phenotyped. Rescue of the *deNimes* phenotype by expression of transgenic *sev* under a heat-shock promoter was performed as described in Ref. ^38^. Vials with wandering larvae and prepupae were subjected to a single 30-minute heat shock at 36°C in a water bath and allowed to develop to the adult stage. *deNimes/Y; hs-sev/+* males were phenotyped.

Genomic DNA was isolated using standard methods from wild-type and *deNimes* flies, and regions of the *sev* gene were amplified. PCR products shown in the Supplementary Fig. 1D were amplified using the following primer combinations (expected product size from the wild-type template in parentheses): s1, sev701 + sev704 (2000 bp); s2, sev701 + sev703 (1653 bp); s3, sev700 + sev704 (1283 bp); s4, sev700 + sev703 (936 bp); a1, sev572 + sev704 (648 bp); a2, sev700 + sev516 (431 bp). Primer sequences are listed in Supplementary Table 2.

### DGRP lines

The DGRP lines were obtained from the Bloomington *Drosophila* Stock Center (BDSC). Flies were aged for ≥2 weeks, and 5–33 individuals per line were phenotyped across 1–6 experiments by staining for Rh5 and Rh6 expression and counterstaining with phalloidin.

### Laboratory *Drosophila* strains

For wild-type controls, we used *yw; Sp/CyO; wt^isoB^* flies (Ref. ^30^), where ‘isoB’ represents an isogenised wild-type third chromosome. We used the following published strains: *melt^Δ1^* (ref. ^42^); “*melt*-”, *Rh6^1^*(Ref. ^30^; “*Rh6-*”), *Rh5^2^* (Ref. ^66^; “*Rh5-*”). The *sev*^14^ (Ref. ^67^) allele was obtained from BDSC stock # 5691, and the *hs-sev* transgene^38^ from BDSC stock # 10546. For this study we generated the following additional fly lines.

***The melt^Δ160^ deletion mutant*** was generated by CRISPR-Cas9-mediated deletion. gRNAs flanking the 214 bp sequence deleted in the DGRP-509 line were designed using the DRSC/TRiP Functional Genomic Resources Center (https://www.flyrnai.org/crispr/) tool (oligo sequences in Supplementary Table 3) and cloned into the BbsI-digested pCFD3-dU6:3gRNA plasmid^68^ (a gift from Simon Bullock (Addgene plasmid # 49410; RRID:Addgene_49410)). gRNA plasmids were co-injected into the *vas-Cas9*(X) stock^69^ together with an oligo (CTTCCTTGATGGCGCGACAGCCGCGCGAAATGACTTTTCTGCCAGCTCTTTTCATTTCA TGTTTTTTTTTTAGCG*CTCACGGAAAACAGAGCCGCCGAATCATTTTCGCAGAATTTCT GGCCGGCCATACGGTGCAGATTCTTGAGTAAA; asterisk indicates location of the intended deletion) to serve as a donor template for the homology-directed repair. The progeny were screened by PCR for deletions in *melt*. Two hits were identified; in both, sequencing revealed a deletion of 160 bp (deleted sequence: ATCCCTTGCCGGAATTTTGCTCATCTAATCAAAAGACCTGCATTCCACGAAGGTCCAAAG AACTCGACGGCAAGTTGGCCAAGAATTTACCGGTGCTATCAGCGGAACCAACAAGCGG

CCAACTGCAAGTGGGCAAACACAAATAGCGTTTGCGGGAAAC) instead of the expected 214 bp (Supplementary Table 1). This 160 bp deletion disrupted a K_50_/Otd motif (TAATCC)^43,70^ by deletion of 4 bp (ATCC) and removed a Sd motif (identified based on the PWM in FlyFactorSurvey^43^ and similarity to the two Sd motifs found in the *Rh5* promoter^71,72^).

***Wild-type and mutant melt reporter constructs*** were based on the *melt1B* enhancer^25^*, a* 2043 bp *melt* fragment (see sequence below) obtained by restriction digestion of a TOPO plasmid containing a ∼4 kb fragment of the *melt* locus^27^ (a gift from David Jukam, NYU) with KpnI and NotI. Point mutations in the Sd motif, K_50_/Otd motif, or a 214 bp deletion containing these motifs were introduced with the QuikChange II Site-Directed Mutagenesis Kit (Agilent) or the Q5 Site-Directed Mutagenesis Kit (NEB) (primer sequences in Supplementary Table 4). Mutations were confirmed by sequencing. Wild-type and the mutant fragments were cloned into a transformation vector^71^ containing an *hsp70* minimal promoter, the transformation marker *mini-white*, the reporter *egfp*, and an *attB* site for genomic integration^73^ into the third-chromosomal landing site J36 (ZH-attP-86Fb)^73^.

Sequence of the *melt1B* enhancer fragment (2043 bp):

ttgcgtcatcgagtatattcaaatatcaccccgtgagccgcctagtcctagaagtcttattgtgggtttaacaaattcttcaaaaatactgattcaaaatacatttcattacctaaaatggatt gtagtttcaaaacatatacatttcgtgattttgaaatgcttaaaccttgagtaatcacaataattgacccaaattgctgtccaacagtccaagaaattcgactgtaatagaaaggcttgacg aactaagatatttagtggctggctttattccgttcctgaactctgccattttgtggagcaatgcaaatagaatctaagcgaagagttatggggccccgaaaattttggcacagcctttgtttc cactctgtctccgaaatctggagaccatgccctcctctaaaccccccccccccttgcgatcatgggcgcggggcttcgtgtgtattaacctgccgcaccggtttcggtttatgttttagcca acgccatcatccattacttatacccgagctcgaaccccttttttcatacatatgtgtgtttctatttctcttttaattttctctcgggcgctcaaagttgctgctgctgctgctgctgccgctgctgttg ctgcgacttgttgctgcttctttatttattttttaaccagtttttctgtttgttgttggtgctgcaacctgagcgcgaatgagccataaataacaacaaccacagcagcccagtcgctgtcttctgtc ccgatttccaggaagcaaatgtctgtccgtccgtccgtccgcttgtcggcttgtccgcccgtcgagaggcccgaacccaaacccaaagcaacccgtcccgtctgcaattcgcgtgca gagccgctttgcccgcctgactggcctctatggattatgcaactcttggttttggtttttggttttcggttttcggctttcagttttcagtttttatttgaaaatttaataacccaggcgcgttggcgtta gcgtctgcttccttgatggcgcgacagccgcgcgaaatgacttttctgccagctcttttcatttcatgttttttttttagcgcccacggatttggccagccagcaaattcccggctgatccgcta agtcgcgtctaatcccttgccggaattttgctcatctaatcaaaagacctgcattccacgaaggtccaaagaactcgacggcaagttggccaagaatttaccggtgctatcagcggaa ccaacaagcggccaactgcaagtgggcaaacacaaatagcgtttgcgggaaaccccacggaaaacagagccgccgaatcattttcgcagaatttctggccggccatacggtgc agattcttgggtaaacaaactagcttgtgcaattaatgtgggccaatcatggactcaatgcgtattagttgaaggttccataaagataattgacccaaatttgtttgccataaagatagttgt tttagctagtatatttcgcttggatttccgatatcacaagttctctggctgggctgcatacatactcgtataacatgaaatatacattccatcagaggctgagcaagggctattctgcctgaga gtatcttagataaacatcgaggcagtgcaacaagtatttggtgtccgcagaatgtctatatttatacaactacagcaacttatatatgcactttcgggacaatcgagaatagaactgctct ggagcaacaagtaaatattttggcgaaacctctatacaaatgcccggcgataagataaaaagtggctgccagaggcgcctgaactttggcaatgggccaattccaaacatggcag tgaaaaaatgcactgactaaaataaacaaatacacccgactaattctcattgtatatagtccatttcaatggagcgtgatttcttttgcaatactaaaatattaagcaataactgcatataa gaatatataaatcggaggcaatttttacaaagatccttattattttggaattaatattttttttagttgctgcagatgaagaatt

### Comparison of our dataset with the ss-screen dataset

We compared the p:y R7 percentages from the ss-screen^16^ with p:y R8 ratios in our Rh5/Rh6 screen for each DGRP line common to both datasets (Supplementary Data 1). Anderson et al. reported sample sizes for all lines in the ss-screen as ≥ 6; therefore, we conservatively assumed n = 6 for their data when calculating statistical significance. Differences between our screen and the ss-screen were assessed using two-sided Welch’s t-tests, with Bonferroni correction for multiple comparisons. As an additional way of prioritising lines for follow-up analysis, we also marked as outliers lines whose difference between the two screens fell outside the lower or upper bounds calculated from the distribution of differences. All calculations are provided in Supplementary Data 1, sheet SD1-8. We identified a few lines with high p:y R7 ratios in the ss-screen that had approximately normal p:y R8 ratios in our dataset. For these lines, we reproduced the ss-screen crosses and, in several cases, observed approximately normal p:y R7 ratios (Supplementary Data 1, sheets SD1-10 and SD1-11). Our comparison used the corrected values (Supplementary Data 1, sheet SD1-8). No conclusions of the original study would be affected by this correction.

## DATA AVAILABILITY

The quantitative data supporting the findings of this study, including individual-retina counts and statistical analyses, are provided in Supplementary Data 1. The genetic variants identified in this study are described in Supplementary Table 1. The publicly available DGRP2 genomic sequence and variant data were previously published and are accessible as described in refs. ^31^ and ^32^. Raw microscopy images are available from the corresponding author upon reasonable request.

## Acknowledgements

We thank Amir Yassin, Nikos Konstantinides, Virginie Courtier-Orgogozo, Abhishek Chatterjee, and Claude Desplan for helpful comments on the manuscript. We thank Robert J. Johnston, David Jukam, Simon Sprecher, Michael Perry and François Rouyer for helpful discussions and exchanges. We are very grateful to Darren Obbard, Clémentine Vasiliauskas and Monique Douard for collecting wild flies used in this work. We are indebted to Steve Britt, Michael Perry and Claude Desplan for generously providing the Rhodopsin antibodies. We warmly thank Béatrice Martin for generous practical assistance throughout the project, and Eric Mocoeur for valuable technical contributions. Fly stocks obtained from the Bloomington *Drosophila* Stock Center (NIH P40OD018537) were used in this study.

## Funding

This work was supported by the following grants to D.V.: an EU FP7 Marie Curie Career Integration Grant (CIG), grant no. 631687 (COLOURHODOPSIN); a Lidex NeuroSaclay, Université Paris-Saclay grant (NaturalNeuro); and an Agence Nationale de la Recherche grant no. ANR-21-CE12-0038 (WILD_EYES). It was also supported by the following grants to J.R.: R00EY023995 and R01EY036459 from the National Eye Institute/NIH.

## Author contributions

D.V. conceived and supervised the study. D.V. and J.R. designed experiments and obtained funding. D.N., S.G., J.B., B.M., F.V., S.L., R.E.C., D.D., F.A., L.T. and D.V. performed experiments. D.N., S.G., J.B., B.M., F.V., F.A., L.T., J.R. and D.V. analysed and interpreted data. D.N., S.G., J.B., B.M., F.V., F.A., L.T. and D.V. developed methods or reagents. D.V. prepared the figures. D.V. and A.B.P. wrote the manuscript, with substantial revisions by J.R. and L.T.

## Competing interests

The authors declare no competing interests.

## Notes

### Competing Interest Statement

The authors have declared no competing interest.

### Summary of Updates

The Introduction was reorganised, and the Discussion substantially revised. The Results were lightly edited to improve clarity. One author was added. Typos were corrected in Supplementary Table 1.

