## Supplementary Figures 1-2 and Supplementary Tables 2-4 for "Natural genetic variation spans the qualitative phenotypic space of a neuronal identity endophenotype"

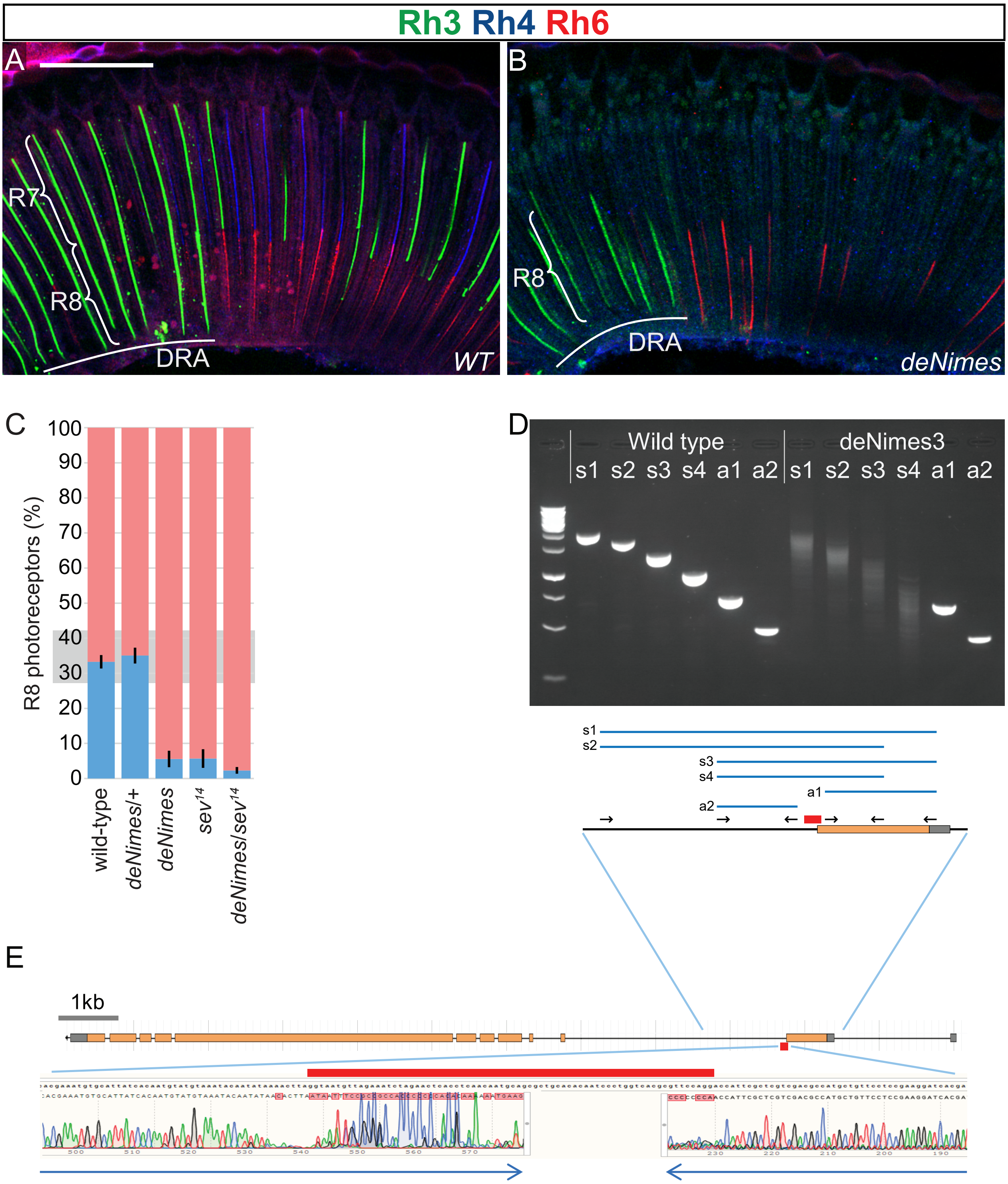


### Supplementary Figure 1

**Supplementary Figure 1. *deNimes* lacks R7 Rhodopsin expression and carries an altered *sevenless* locus.**

A–B. Side views of ommatidia near the rim of the adult retina, including the Dorsal Rim Area (DRA, underlined). A. In wild-type retina, Rh3 (green) and Rh4 (blue) are expressed in R7, whereas Rh6 (red) is expressed in R8 in the main retina. In DRA ommatidia, both R7 and R8 express Rh3. B. In the *deNimes* retina, R8 photoreceptors still express Rh3 (in the DRA) and Rh6, but no Rh3 or Rh4 expression is detected in the layer above R8. Braces indicate R7 and R8 layers. Scale bar, 50 µm.

C. Quantified Rh5/Rh6 expression phenotypes of wild-type, *deNimes* heterozygotes and homozygotes, *sevenless* (*sev*) mutants, and *deNimes*/*sev* trans-heterozygotes, showing recessive inheritance and failure to complement *sev*. Each vertical bar represents one genotype, with the fraction of Rh6-expressing R8s in red and Rh5-expressing R8s in blue. Error bars indicate SD of pR8 frequency for each line. The shaded band centred at ~33% indicates the typical range of p:y ratios observed in laboratory wild-type strains.

D. Agarose gel of PCR products from amplification of the *sev* genomic locus from wild-type and *deNimes* flies. A schematic of the *sev* genomic region is shown below the gel (a detail of the map in (E)), indicating the approximate locations of the primers used (arrows) and the PCR products (blue) labelled as on the gel. s1–s4, PCR products amplified with different primer pairs spanning the variant region (red, see also E). a1, a2, PCR products amplified with primer pairs positioned adjacent to, but on the same side of, the variant region. Spanning primer pairs produce smeared products from the *deNimes* template that are similar in size to the corresponding wild-type bands, but migrate slightly faster, whereas adjacent primer pairs produce discrete bands from both templates.

E. Top, *sev* locus (transcribed right to left) showing exon–intron structure and the protein-coding sequence (orange). Bottom, examples of sequence traces from both directions (blue arrows) of the *deNimes* s1 PCR product shown in (D), aligned to the wild-type *sev* sequence. The readable sequence in the traces ends abruptly without overlap. Red bars above the sequence (bottom) and below the map (top) indicate the 72 bp of wild-type sequence affected in *deNimes* flies.

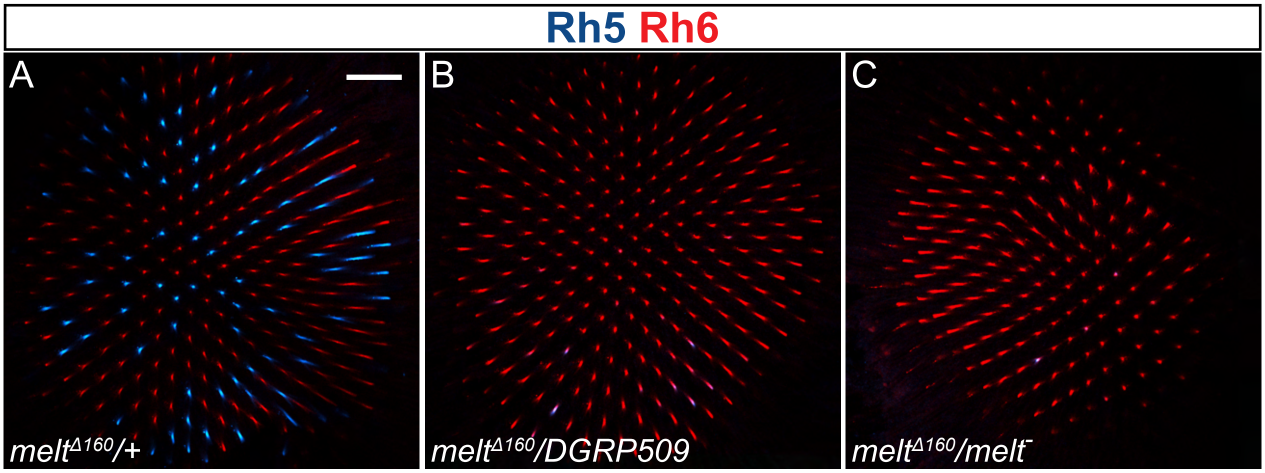


Supplementary Figure 2

**Supplementary Figure 2. Complementation tests for a CRISPR-generated 160 bp deletion in the *melt* locus.**

A–C. A CRISPR-generated 160 bp deletion in the *melt* locus (see Fig. 10G and text for details) is recessive (A) and fails to complement the genetic variant causing the DGRP-509 phenotype (B) or the null melt mutation (C). All micrographs are optical sections through central wholemount retinas stained with antibodies to reveal Rh5 (blue) and Rh6 (red). Scale bar, 25 µm.

Supplementary Table 2. Primers used for PCR amplification of the *sevenless* locus.

| Primer | Direction (relative to the genome) | Sequence |
|---|---|---|
| sev700 | forward | AAGCCGAGCCGAGACGAGGAGGTAA |
| sev701 | forward | CCGATTACACACTCTGCGATA |
| sev703 | reverse | CCTGCTCATTCTACTCTCGATCT |
| sev704 | reverse | AGTGCCTCGATGACTATGTTTTG |
| sev516 | reverse | GACATTCGGGTGCGAAATGT |
| sev572 | forward | CCAGGACCATTCGCTCGT |

Supplementary Table 3. Oligos for gRNAs for CRISPR-Cas9-mediated *melt* genomic deletion.

| | **Sense** | **Antisense** |
|---|---|---|
| 5' gRNA 1 | GTCGAGTCGCGTCTAATCCCTTGC | AAACGCAAGGGATTAGACGCGACT |
| 5' gRNA 2 | GTCGACGCGACTTAGCGGATCAGC | AAACGCTGATCCGCTAAGTCGCGT |
| 3' gRNA 1 | GTCGTCGGCGGCTCTGTTTTCCGT | AAACACGGAAAACAGAGCCGCCGA |
| 3' gRNA 2 | GTCGGCCATACGGTGCAGATTCT | AAACAGAATCTGCACCGTATGGCC |

Supplementary Table 4. Primers used for mutating the *melt1B-GFP* reporter (mutant nucleotides are indicated in lowercase).

| **Mutation** | **Forward Primer** | **Reverse Primer** |
|---|---|---|
| 214 bp deletion | CCCACGGAAAACAGAGCC | CGCTAAAAAAAAAACATGAAATGAAAAGAG |
| $K_{50}$/Otd | AGTCGCGTCT**cgc**CCCTTGCCGG | TAGCGGATCAGCCGGGAA |
| Sd | AAAGACCTGC**caaaa**ACGAAGGTCC | TGATTAGATGAGCAAAATTCC |
